# Vitamin D modulates cortical transcriptome and behavioral phenotypes in an *Mecp2* heterozygous Rett syndrome mouse model

**DOI:** 10.1101/2021.06.30.450587

**Authors:** Mayara C. Ribeiro, Jessica L. MacDonald

**Affiliations:** Department of Biology, Program in Neuroscience, Syracuse University, Syracuse NY

## Abstract

Rett syndrome (RTT) is an X-linked neurological disorder caused by mutations in the transcriptional regulator *MECP2*. *Mecp2* loss-of-function leads to the disruption of many cellular pathways, including aberrant activation of the NF-κB pathway. Genetically attenuating the NF-κB pathway in *Mecp2*-null mice ameliorates hallmark phenotypes of RTT, including reduced dendritic complexity, raising the question of whether NF-κB pathway inhibitors could provide a therapeutic avenue for RTT. Vitamin D is a known inhibitor of NF-κB signaling; further, vitamin D deficiency is prevalent in RTT patients and male *Mecp2*-null mice. We previously demonstrated that vitamin D rescues the aberrant NF-κB activity and reduced neurite outgrowth of *Mecp2*-knockdown cortical neurons *in vitro*, and that dietary vitamin D supplementation rescues decreased dendritic complexity and soma size of neocortical projection neurons in both male hemizygous *Mecp2*-null and female heterozygous mice *in vivo*. Here, we have identified over 200 genes whose dysregulated expression in the *Mecp2*+/- cortex is modulated by dietary vitamin D. Genes normalized with vitamin D supplementation are involved in dendritic complexity, synapses, and neuronal projections, suggesting that the rescue of their expression could underpin the rescue of neuronal morphology. Further, motor and anxiety-like behavioral phenotypes in *Mecp2*+/- mice correlate with circulating vitamin D levels, and there is a disruption in the homeostasis of the vitamin D synthesis pathway in *Mecp2*+/- mice. Thus, our data indicate that vitamin D modulates RTT pathology and its supplementation could provide a simple and cost-effective partial therapeutic for RTT.

## Introduction

Rett syndrome (RTT) is a complex neurological disorder caused by mutations in the transcriptional regulator *MECP2* (Amir et al., 1999). *MECP2* (*Mecp2* in mouse) is located on the X-chromosome and RTT patients are predominantly females who are heterozygous for *MECP2* mutations. Due to X-inactivation, they display cellular mosaicism with approximately half of their cells expressing the mutant allele of *MECP2* and approximately half expressing the functioning allele. Female RTT patients go through a period of regression after the first 6-18 months of life, including the loss of motor skills and purposeful use of their hands. Additional symptoms include deceleration of head growth, ataxia, breathing abnormalities, and the onset of seizures and autistic behaviors (Chahrour and Zoghbi, 2007). *Mecp2*-mutant mouse models have been widely utilized to investigate the role of MeCP2 in the CNS and the progression of RTT phenotypes. These models recapitulate many aspects of RTT, including behavioral phenotypes such motor deficits (Samaco et al., 2013; Vogel Ciernia et al., 2018, 2017), impaired cognition (Pelka et al., 2006), social dysfunction (Kerr et al., 2008; Moretti et al., 2005; Pearson et al., 2012; Samaco et al., 2013; Schaevitz et al., 2010), and altered anxiety-like behavior (Samaco et al., 2013; Vogel Ciernia et al., 2017). *Mecp2*-mutant mouse models have also provided crucial insight into the mechanisms of action of MeCP2, its transcriptional targets, the proteome disruptions that occur with its loss-of-function, and the resulting phenotypes.

One of the many disrupted cellular pathways identified in the CNS of *Mecp2*-null mice is aberrant NF-κB signaling (Kishi et al., 2016). Importantly, genetic attenuation of the NF-κB pathway, by crossing *Mecp2* deficient mice with mice that are heterozygous for the *Nfkb1* subunit of NF-κB, is sufficient to extend lifespan in male *Mecp2*-null mice and rescue dendritic complexity of projection neurons within the superficial layers of the neocortex (Kishi et al., 2016). Further, inhibiting the NF-κB pathway via inhibitors of Gsk3b reduces proinflammatory cytokine expression and rescues dendritic and spine deficits in *Mecp2*-null neurons (Jorge-Torres et al., 2018). Thus, NF-κB inhibition could provide a therapeutic approach for RTT.

Vitamin D is a known inhibitor of the NF-κB pathway (Al-Rasheed et al., 2015; Y. Chen et al., 2013; D’Ambrosio et al., 1998; Giarratana et al., 2004; Penna et al., 2009; Sun et al., 2006), and it has been shown to decrease total NF-κB protein levels and reverse cognitive deficits induced by high-fat diet in rats (Hajiluian et al., 2017), and to decrease proinflammatory cytokines induced by NF-κB (Guillot et al., 2010; Harant et al., 1997; Mokhtari-Zaer et al., 2020). Vitamin D can also regulate multiple other signaling pathways that are disrupted in RTT, including BDNF, IGF1, and the P13K/AKT pathway (reviwed in Marballi and MacDonald, 2021), suggesting it could improve RTT phenotypes through modulation of multiple signaling pathways. Notably, vitamin D deficiency is prevalent in RTT patients (Motil et al., 2011; Sarajlija et al., 2013) and *Mecp2*-null mice (Ribeiro et al., 2020). This led us to investigate whether vitamin D supplementation can rescue the aberrant NF-κB pathway activation and improve phenotypes in *Mecp2*-mutant mice, and to investigate the gene expression changes that could underpin this rescue.

Previously, we found that adding the activated form of vitamin D to *Mecp2*-knockdown cortical neurons *in vitro* rescues their aberrant NF-κB activation and reduced neurite outgrowth. Further, we demonstrated that vitamin D dietary supplementation beginning at 4 weeks of age (early symptomatic) rescues the reduced dendritic complexity and soma size of neocortical superficial layer neurons in male *Mecp2*-null mice at 8 weeks of age, and significantly (∼20%) improves their reduced lifespan (Ribeiro et al., 2020). We extended this work into the more clinically relevant female heterozygous model (Ribeiro and MacDonald, 2020), identifying a partial rescue of the reduced dendritic complexity and soma size phenotypes at 5 months of age, with sex-specific differences in dose (Ribeiro et al., 2020). These results provide evidence that vitamin D supplementation could be a partial therapeutic option for RTT.

In this study, we investigated whether dietary vitamin D supplementation can rescue the expression of genes that are dysregulated within the neocortex of *Mecp2*+/- mice, and whether vitamin D deficiency further exacerbates transcriptome disruptions in these mice. We found that dietary vitamin D modification has a profound impact on the transcriptome of the neocortex. We identified more than 200 differentially expressed genes whose expression is normalized with vitamin D supplementation, many of which are associated with neuronal morphology. Dietary vitamin D deficiency exacerbated the dysregulation of many of these genes in the *Mecp2*+/- cortex, but, strikingly, it normalized the expression of many other dysregulated genes, similar to the effect of supplementation. We further found that motor deficits and anxiety-like behavior in the open field are exacerbated in *Mecp2*+/- mice with insufficient circulating vitamin D, while *Mecp2*+/- mice with either sufficient or deficient circulating vitamin D show behavioral improvements. We identified a disruption in the homeostasis of the vitamin D synthesis pathway in *Mecp2*+/- mice, which could contribute to the enhanced sensitivity of *Mecp2*+/- mice to vitamin D levels, with a potential compensation with extremely low or high levels of vitamin D. Thus, our data indicate that vitamin D can modulate RTT pathology, and its supplementation could provide a simple and cost-effective partial therapeutic for RTT.

## Materials and Methods

### Animals

All animal experimental protocols were approved by the Syracuse University Institutional Animal Care and Use Committee and adhere to NIH ARRIVE guidelines. Mice were group housed at a maximum of five mice per cage on a 12/12 h light/dark cycle and were given food and water ad libitum. Female *Mecp2* heterozygous mice were purchased from The Jackson Laboratory (B6.129P2(C)-Mecp2tm1.1Bird/J; RRID:IMSR_JAX:003890), and were maintained on a C57BL/6 background. Genotypes were determined by PCR on genomic DNA as follow: *Mecp2* mutant mice, forward primer oIMR1436 5’ -GGT AAA GAC CCA TGT GAC CC - 3’; reverse primer oIMR1437 5’ - TCC ACC TAG CCT GCC TGT AC - 3’; reverse primer oIMR1438 5’ - GGC TTGCCACATGACAA - 3’. Mice were weighed weekly by an investigator blinded to genotype and chow concentration.

### Vitamin D supplementation and serum measurements

Custom chow obtained from Bio-Serv was based on the AIN-93G Rodent Diet, varying only in Vitamin D3 concentration. Female *Mecp2*+/+ and *Mecp2*+/- littermates were each weaned together at 4 weeks of age and placed on chow containing 1 IU/g (control diet), 10 IU/g (supplemented diet) or 0.1 IU/g (deficient diet) vitamin D in rotating order based on date of birth.

Serum was collected from *Mecp2*+/+ and *Mecp2*+/- females on all chows at 7 months of age, after the completion of the behavioral assays. Total serum 25(OH)D levels were measured via mass spectrometry by ZRT Laboratories (Beaverton, OR).

### RNA-sequencing

Total RNA was extracted from the cortices of 7-month-old mice using TRIzol reagent (Invitrogen) and RNeasy Mini Kit (Qiagen) (Untergasser, 2008). The samples consisted of 3 *Mecp2*+/+ and 3 *Mecp2*+/- on each diet (1IU/g, 10IU and 0.1IU/g vitamin D). Libraries were made using QuantSeq 3’ mRNA-Seq Library Prep Kit FWD for Illumina (Lexogen) according to the manufacturer’s instructions. The libraries were sequenced on an Illumina NextSeq 500 next-generation sequencing instrument using the NextSeq 500/550 High Output kit v2.5 (75 cycles) (Illumina) to generate approximately 22M reads per sample.

#### Analysis

RNA-sequencing analysis was performed with the PartekFlow software. In brief, adapters and bases were trimmed, followed by alignment with STAR 2.7.3a index (mus musculus mm10 assembly, whole genome aligner index) and quantification to model (Gencode M25). Normalization to remove unwanted variation (RUV) was performed with R (Risso et al., 2014). Gene specific analysis (GSA) and heatmaps were done on PartekFlow. Genes were deemed differentially expressed based on their p-value (<0.05) and false discovery rate (FDR<0.1). Volcano plots were made in GraphPad Prism. Upset plots were created with the web-version of UpsetR (Lex et al., 2014), while gene ontology (GO) analysis and transcription factors enrichment were performed on g:Profiler (Raudvere et al., 2019).

### Behavioral analyses

Behavior was assessed at 3, 5 and 7 months of age. All behavioral tests (described in detail below) were run during the light cycle. The mice were acclimated to the behavior room for at least 30 min prior to testing.

#### Accelerating Rotarod

The rotarod apparatus (Ugo Basile) was set to accelerate from 5 to 50 rpm in an interval of 5 min. Each mouse was tested three times per day, with one hour between trials, for three consecutive days. Each trial entailed the placement of the mouse on the stationary apparatus and given 5 sec to acclimate to its movement before it was accelerated. In cases where the mouse fell from the beam before 30 sec had elapsed, the test was reset, and the mouse was placed back on the apparatus; this was repeated for a total of three times. The latency to trial end, which indicates the fall of the mouse to the floor of the rotarod, was recorded for each mouse. The trial was manually ended if the mouse clutched the rotating rod without walking for five consecutive turns (Vogel Ciernia et al., 2017).

#### Open field exploration

Each mouse was placed in the center of a 40 cm x 40 cm open field maze with a photo-beam recording system (SD Instrument) for 30 min. Total distance travelled and time/distance spent in the center of the maze (10 cm x 10 cm, equivalent to ¼ the area of the apparatus) was quantified with San Diego’s PAS Reporter software.

#### Elevated plus maze

Each mouse was placed in the center of a plus-shaped maze (arms: 35 cm x 5 cm; wall: 20 cm in height) elevated 63.5 cm off the ground for a total period of 5 min. Travel distance, time spent in each arm, number of entries in each arm and time spent with the nose over the edge of the open arms were quantified with the Ethovision XT (Noldus) software.

#### Social approach

The social approach apparatus consisted of three chambers (40 cm x 20 cm) connected by sliding doors. Each mouse was placed in the closed center chamber for 5 min. After, the doors were removed, and the mouse was free to explore all three chambers for another 10 min to acclimate to the apparatus. After acclimation, the mouse was returned to the enclosed center chamber and one wire pencil cup was placed in the right chamber and another identical cup was placed in the left chamber. One of the cups contained an age and sex matched mouse (novel mouse) while the other cup remained empty (novel object) (Vogel Ciernia et al., 2017). After the apparatus was set, the doors were open once again and the mouse was free to explore the chambers for 10 min. The Ethovision XT (Noldus) software was used to calculate total distance travelled, time spent in each chamber, number of entries to each chamber, and time sniffing the novel mouse/object.

### Quantitative reverse transcription PCR (RT-qPCR)

RNA was extracted using TRIzol (Invitrogen), and cDNA was synthesized using qScript cDNA SuperMix (Quanta Biosciences). RT-qPCR was performed on a CFX Connect Real-Time System (Bio-Rad Laboratories) according to the manufacturer’s instructions. Primer pairs were designed in intron-spanning regions or exon-exon junctions as to not amplify genomic DNA. The primer pairs were as follow:

*CYP2R1*: forward 5’ - TGTATGGCGAGATTTTCAGTTTAG - 3’; reverse 5’ - CAAAGGAAGGCATGGTCTATCT - 3’.

*CYP27A1*: forward 5’ - CATGGATCAGTGGAAGGACC - 3’; reverse 5’ - GCCTCTGTTTCAAAGCCTGA - 3’ (Zeisel A, Yitzhaky A, Bossel Ben-Moshe N, 2013).

*CYP27B1*: forward 5’ - AGTGTTGAGATTGTACCCTGTG - 3’; reverse 5’ TAGGGAGACTAGCGTATCTTGG - 3’.

*CYP24A1*: forward 5’ - AACTGTACGCTGCTGTCACG - 3’; reverse 5’ - TCTCTGTTGCACTTGGGGAT - 3’ (Zeisel A, Yitzhaky A, Bossel Ben-Moshe N, 2013).

We used PerfeCTa SYBR Green FastMix (Quanta Biosciences) Master mix, and each PCR consisted of 1X LightCycler FastStart DNA Master SYBR Green I mixture, 0.2 μM primers, and cDNA. We used the mean of *Gapdh* and *S16* expressions as the reference gene. Each sample was run in triplicate and averaged. The relative quantification analysis was performed as follow: ΔCq = Cq of gene of interest - geometric mean of Cq of reference genes; ΔΔCq = ΔCq - mean of ΔCq of wild-type samples; fold change = 2^-ΔΔCq^. Melt curve analysis was also performed to verify specificity of the amplicons.

### Statistical analysis

GraphPad Prism 8.0 (GraphPad Software) was used to carry out the statistical analyses. No statistical method was used to predetermine sample sizes, but our sample sizes are similar to those generally employed in the field. Our statistical tests consisted of two-tailed t test, or two-way ANOVA with Tukey’s multiple comparison. Data distribution was handled as if normal, but this was not formally tested (since potential differences in results would be minor). All data shown represent mean ±SEM. Sample size and statistical test are specified in each figure legend.

## Results

### Vitamin D supplementation normalizes the expression of dysregulated genes in *Mecp2*+/- cortex, many of which are associated with neuronal morphology

Having identified morphological rescue of cortical neurons with vitamin D supplementation, we investigated the underlying molecular mechanisms. Female heterozygous *Mecp2*+/- mice and wildtype (*Mecp2*+/+) littermates were placed on control (1 IU/g vitamin D) or supplemented (10 IU/g vitamin D) chow at one month of age, before the onset of overt behavioral phenotypes. We chose 10 IU/g vitamin D chow as it was the dose that improved neuronal morphology phenotypes of *Mecp2*+/- mice most effectively (Ribeiro et al., 2020). RNA-sequencing was performed on the total neocortex at 7 months of age, when phenotypes are readily apparent. We first confirmed that dietary vitamin D supplementation modifies gene expression within the neocortex. We identified over 3,000 differentially expressed genes (DEG) in the cortex of *Mecp2*+/+ mice on supplemented chow compared to *Mecp2*+/+ on the control diet (2,680 upregulated and 1,177 downregulated; p < 0.05, FDR < 0.1), indicating that many genes within the cortex are responsive to vitamin D supplementation (Fig. 1A). These vitamin D responsive genes are enriched in a broad range GO Molecular Function categories, including chromatin binding, RNA binding, and transcription coregulator activity. Further, 1,350 of these genes contain the vitamin D receptor (VDR) transcription factor binding motif (GGGKNARNRRGGWSA), indicating that they could be directly regulated by vitamin D. Thus, the cortical gene expression changes found with vitamin D supplementation likely result from both direct and indirect (or secondary) regulation.

**Figure 1.**
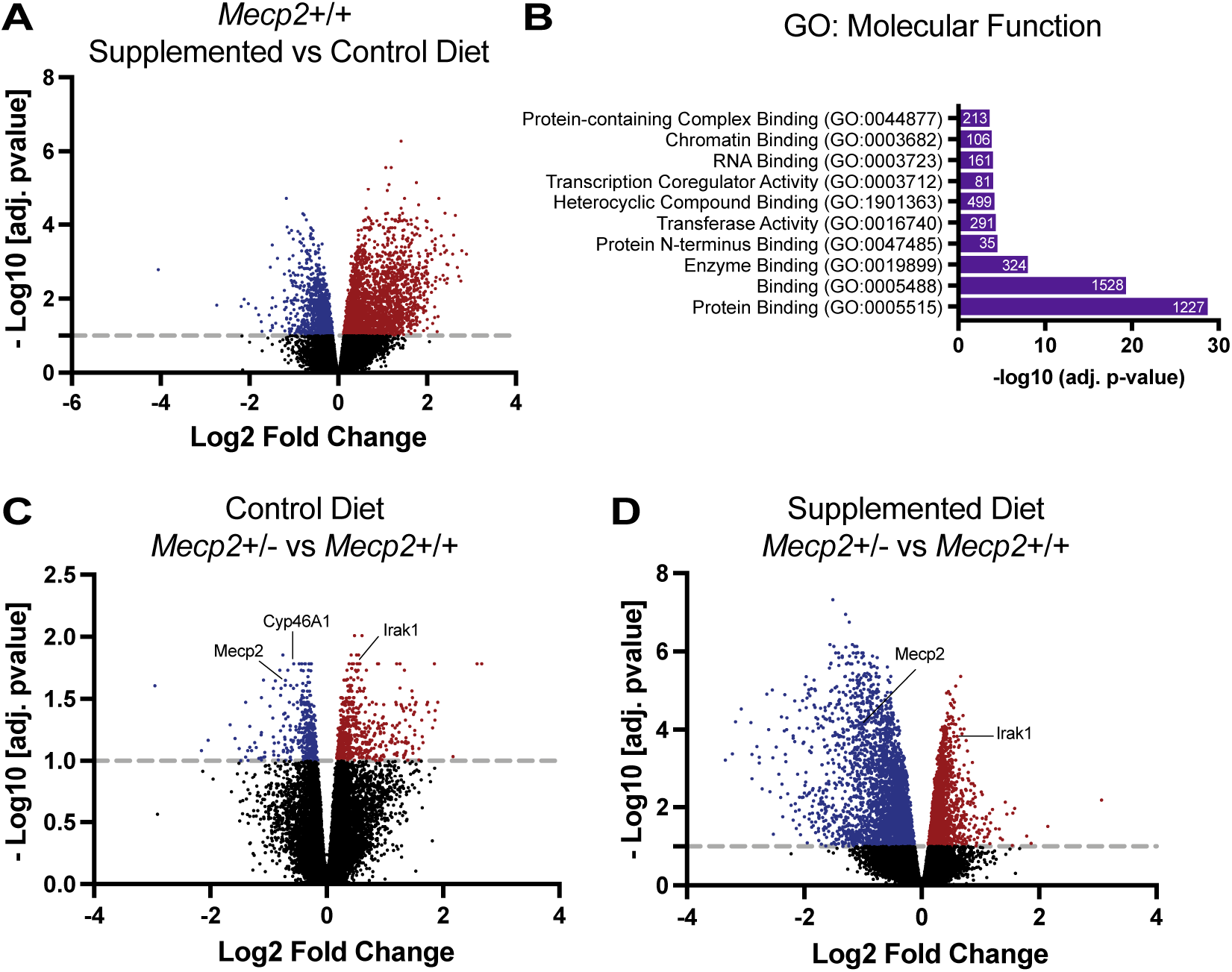
Vitamin D supplementation leads to extensive transcriptome changes in the cortex of *Mecp2*+/+ and *Mecp2*+/- mice. (A) Volcano plot depicting the changes in gene expression between *Mecp2*+/+ cortex on the vitamin D supplemented diet relative to the control diet. Vitamin D supplementation leads to the upregulation of 2,680 genes and the downregulation of 1,177 genes. (B) Gene Ontology analysis of Cellular Components enriched in differentially expressed genes (DEGs) between the cortex of *Mecp2*+/+ mice on the control versus supplemented vitamin D diet. The 10 most significant categories (out of 86 total) are shown. (C) Volcano plot displaying the DEGs between *Mecp2*+/- and *Mecp2*+/+ cortex on the control diet. More DEGs are upregulated in *Mecp2*+/- cortex than downregulated. *Cyp46a1* is downregulated in the cortex of *Mecp2*+/- animals, while *Irak1* expression is upregulated, confirming previously reported data. (D) Volcano plot depicting the DEGs between *Mecp2*+/- and *Mecp2*+/+ mice on the supplemented diet. Vitamin D supplementation results in increased downregulation of genes in the cortex of *Mecp2*+/- mice when compared to their wildtype littermates. Vitamin D treatment normalizes the expression of *Cyp46a1* whereas expression of *Irak1* remains upregulated. A, C, D: Gray dotted line indicates FDR = 0.1. Red indicates upregulated genes and blue indicates downregulated genes. DEG = differentially expressed gene.

We next asked whether the expression of genes that are differentially expressed in the cortex of *Mecp2*+/- mice, relative to wildtype, on the control chow are rescued or normalized with vitamin D supplementation. We found 989 DEGs (p < 0.05, FDR < 0.1) in the cortex of *Mecp2*+/- mice on the control diet when compared to their wild-type littermates (Fig. 1C) but 5,233 DEGs between *Mecp2*+/- and *Mecp2*+/+ treated with the supplemented chow (Fig. 1D). We confirmed that *Mecp2* expression is reduced in *Mecp2*+/- cortex on both control and supplemented chow and expression of *Irak1*, the kinase upstream of the NF-κB pathway, is increased in the cortex of *Mecp2*+/- mice on both control and supplemented diets (Fig. 1C-D).

We hypothesized that genes whose expression is normalized with vitamin D supplementation underpin the morphological rescue that occurs in *Mecp2*-mutant neurons with vitamin D supplementation (Ribeiro et al., 2020). Thus, we considered to be rescued the genes that are differentially expressed in the cortex of *Mecp2*+/- females on control chow, when compared to their wildtype littermates, but that are not differentially expressed in *Mecp2*+/- treated with the supplemented diet (Fig. 2A). We identified 283 rescued DEGs (Supplemental Table 1), the majority of which are downregulated in the *Mecp2*+/- mice on control diet (Fig. 2B). According to GO analysis of cellular components, these rescued DEGs are significantly enriched for neuronal structure, including dendritic tree, synapse, and neuron projection (Fig. 2C; Supplemental Table 2). Deficits in neuronal morphology are a hallmark phenotype in both RTT model mice (Belichenko et al., 2009; Fukuda et al., 2005; Kishi and Macklis, 2004; Rietveld et al., 2015; Stuss et al., 2012; Tropea et al., 2009) and RTT patients (Armstrong et al., 1995, 1999; Armstrong, 2002; Jellinger et al., 1988). Our previous work demonstrated that vitamin D supplementation rescues neurite outgrowth of cortical neurons *in vitro* and dendritic complexity and soma area in male and female mice *in vivo* (Ribeiro et al., 2020). Thus, it is likely that the rescued DEGs identified here are involved in the phenotypic amelioration of cortical neuronal morphology of *Mecp2*-mutant mice after vitamin D supplementation.

**Figure 2.**
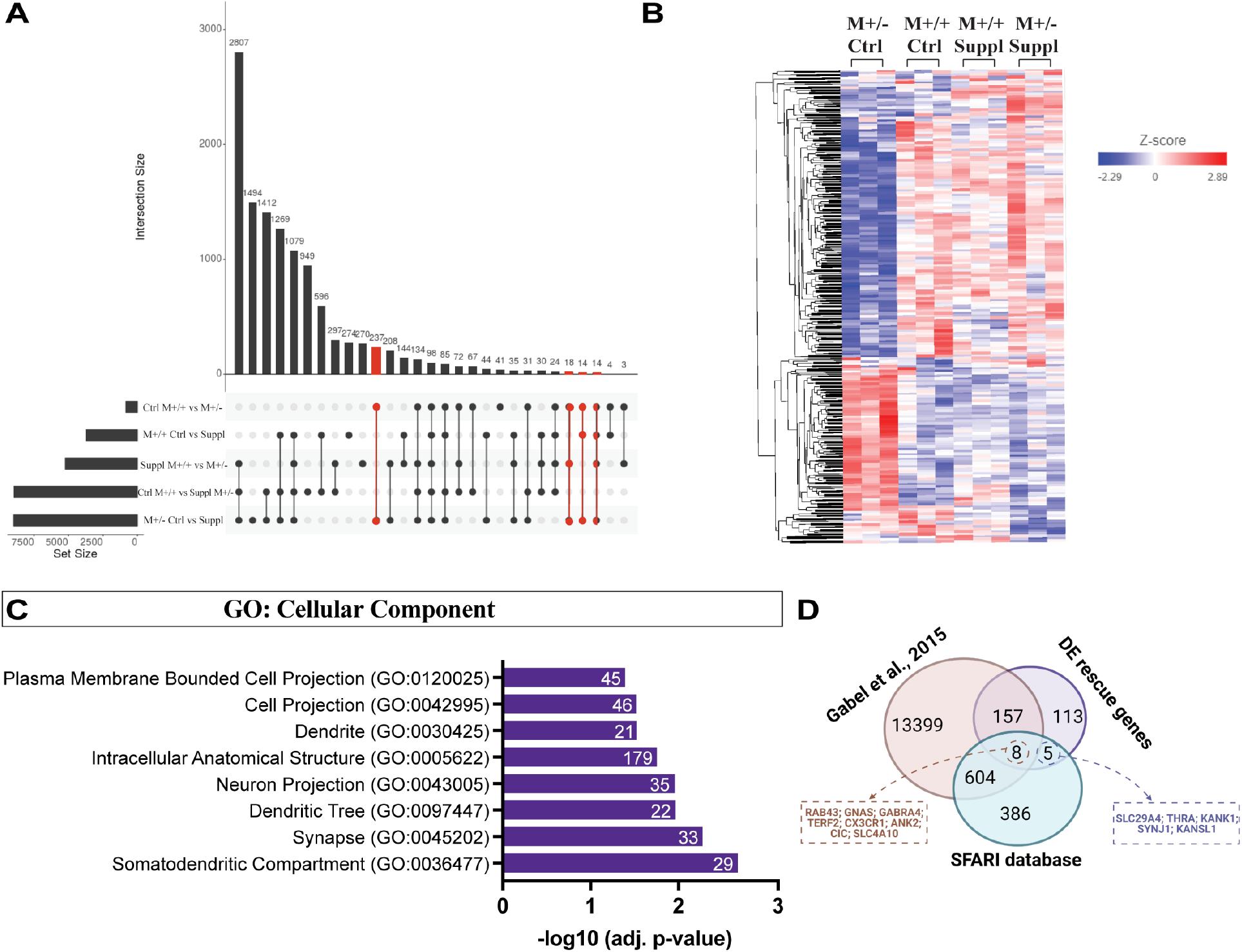
Vitamin D supplementation normalizes the expression of dysregulated genes in the cortex of *Mecp2*+/- mice. (A) Upset plot illustrating the number of DEGs when comparing across genotypes and diets. Red indicates groups that were selected as meeting the criteria for DEGs that are normalized in the cortex of *Mecp2*+/- mice upon vitamin D supplementation. (B) Heat map showing the pattern of expression of the DEGs in the cortex of *Mecp2*+/- (M+/-) mice on control diet (Ctrl) that were normalized with vitamin D supplementation (Suppl), such that they are no longer differentially expressed from *Mecp2*+/+ cortex (M+/+). (C) GO of cellular components analysis of the rescued DEGs, indicating that they are significantly enriched for genes involved in neuronal morphology. (D) Venn diagram illustrating the intersection between the rescued DEGs, genes previously identified as being dysregulated in *Mecp2*-mutant mice (Gabel et al, 2015), and genes associated with ASD (SFARI database).

Of the 283 rescued DEGs, 165 have been previously found to be altered in *Mecp2*-mutant mice (Gabel et al., 2015), supporting the likely importance of these rescued DEGs to RTT pathophysiology. Importantly, we have identified 118 DEGs that have not previously been identified in the compilation of altered RTT genes (Fig. 2D). Further, 8 of the previously RTT-associated rescued DEGs and 5 of the novel RTT-associated rescued DEGs (Fig. 2D) are associated with autism spectrum disorders (ASD), according to the SFARI database, a curated catalogue of genes associated with ASD (Abrahams et al., 2013). Thus, vitamin D might regulate the cortical expression of genes associated with other neurodevelopmental disorders with overlapping pathology with RTT. Notably, 74 of the rescued DEGs contain the vitamin D receptor (VDR) transcription factor binding motif, indicating that they could be directly regulated by vitamin D. Thus, vitamin D should be further evaluated for its potential broad therapeutic benefits in neocortical disruptions.

### Vitamin D deficiency leads to extensive transcriptome changes

To investigate whether vitamin D deficiency further exacerbates the transcriptional dysregulation found in the *Mecp2*+/- cortex, and/or leads to unique disruptions, we additionally placed *Mecp2*+/- mice and littermate controls (*Mecp2*+/+) on a vitamin D deficient diet (0.1 IU/g vitamin D) at 4 weeks of age and performed RNA-seq on the total neocortex at 7 months of age, in parallel with the control and supplemented diets outlined above. Vitamin D deficiency causes extensive transcriptome changes in the neocortex; *Mecp2*+/+ animals on the vitamin D deficient diet display 10,500 DEGs compared to *Mecp2*+/+ on control diet (Fig. 3A). Further, there are 9,671 DEGs between *Mecp2*+/- and *Mecp2*+/+ on the deficient diet (Fig. 3B). The change in gene expression between *Mecp2*+/- mice and their wildtype littermates is far higher on the deficient diet than on control or supplemented diets, highlighting the remarkable impact of vitamin D deficiency on gene expression within the neocortex.

**Figure 3.**
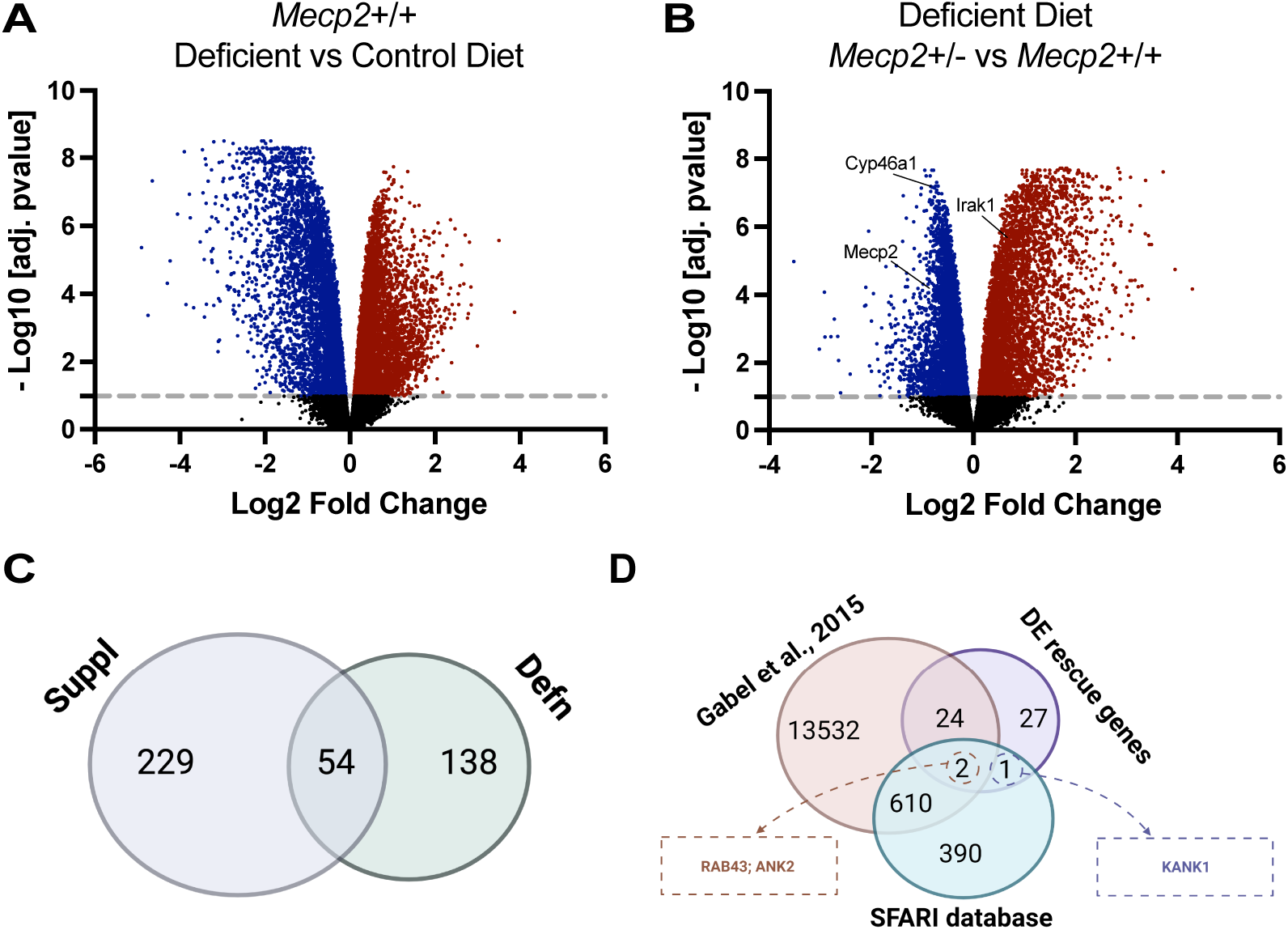
Vitamin D deficiency leads to extensive transcriptome changes in the cortex of *Mecp2*+/+ and *Mecp2*+/- mice. (A) Volcano plot depicting the DEGs between *Mecp2*+/+ cortex on the vitamin D deficient versus control diet. Vitamin D deficiency leads to more downregulation than upregulation of cortical genes. (B) Volcano plot illustrating the DEGs between *Mecp2*+/- and *Mecp2*+/+ cortex on the vitamin D deficient diet. The expression of *Cyp46a1* and *Irak1* remain downregulated and upregulated, respectively, in the cortex of *Mecp2*+/- mice exposed to vitamin D deficient diet, relative to wildtype (*Mecp2*+/+) littermates. (C) Venn diagram illustrating the overlap between the rescued DEGs with vitamin D supplementation and with vitamin D deficiency. (D) Venn diagram illustrating the overlap between the DEGs normalized with vitamin D deficiency, genes previously identified as being dysregulated in *Mecp2*-mutant mice (Gabel et al, 2015), and genes associated with ASD (SFARI database). DEG = differentially expressed genes.

When following the same requisites that we used to select rescued DEGs resulting from vitamin D supplementation, we surprisingly find 192 genes that are differentially expressed between *Mecp2*+/- and *Mecp2*+/+ on control diet that are rescued in *Mecp2*+/- on the deficient diet (Fig. 3C). It should be noted that this list of DEGs rescued by vitamin D deficiency are not enriched for any GO cellular components terms. In addition, none of the DEGs rescued by vitamin D deficiency are enriched for the VDR motif, which is indicative of secondary effects and not direct vitamin D regulation. Interestingly, when comparing the DEGs that were rescued with vitamin D supplementation and the DEGs normalized with its deficiency, we find an overlap of 54 genes (Fig. 3D; Supplemental Table 3). Twenty six of these normalized DEGs have been previously associated with RTT (Gabel et al., 2015) and 3 are found in the SFARI database (Fig. 3.6E). This suggests that the expression of these 54 genes that are dysregulated in the *Mecp2*+/- cortex under control conditions can be normalized with vitamin D modulation, regardless of whether it is supplementation or restriction.

### Vitamin D dietary supplementation ameliorates disrupted motor coordination of *Mecp2*+/- mice and improves anxiety-like behavior in the open-field test

We next investigated whether these alterations in cortical gene expression with vitamin D modulation are correlated with changes in behavior in *Mecp2*+/- mice and wildtype littermates. We placed the animals on control (1 IU/g vitamin D), supplemented (10 IU/g vitamin D), or deficient (0.1 IU/g vitamin D) diets starting at 4 weeks of age, as outlined for the RNA-seq experiments. We performed a longitudinal experiment to evaluate phenotypic progression and longitudinal decline displayed by the *Mecp2*+/- mice, assessing their behavior at early symptomatic (3 months), symptomatic (5 months), and late symptomatic (7 months) stages. We first assessed vitamin D supplementation, hypothesizing that it would improve *Mecp2*+/- phenotypes.

*Mecp2+/-* mice on both control and supplemented vitamin D diets display lower latency to trial end than their *Mecp2+/+* littermates on an accelerating rotarod (Fig. 4A), indicating impaired motor coordination (Deacon, 2013). At 5 months of age, however, the *Mecp2+/-* females on supplemented diet display a significant improvement in motor coordination when compared to *Mecp2+/-* mice on control diet, although this improvement does not persist at 7 months (Fig. 4A). This suggests that dietary vitamin D supplementation delays the decline in motor coordination but does not fully prevent it. Vitamin D supplementation does not significantly alter the total distance travelled (Fig. 4B) or the average speed of locomotion in the open-field test at any of the ages examined (data not shown), however, suggesting that it does not rescue overall locomotory behavior phenotypes of *Mecp2* heterozygous female mice.

**Figure 4.**
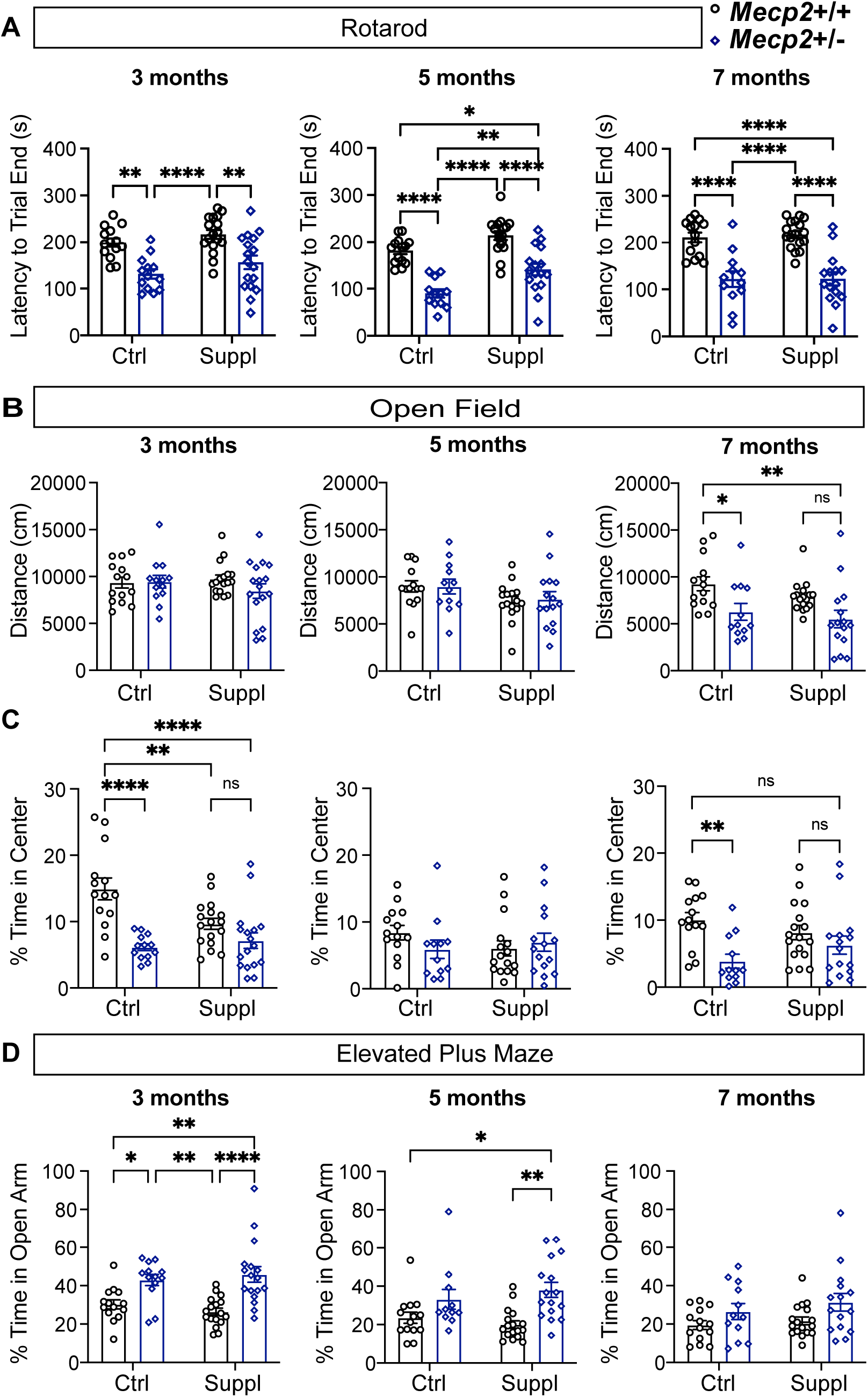
Vitamin D supplementation mitigates some behavioral deficits of *Mecp2*+/- mice. (A) In the rotarod test, *Mecp2*+/- mice (blue) on the control (Ctrl) and vitamin D supplemented (Suppl) diet have a shorter latency to trial end than their wildtype littermates (*Mecp2*+/+; black) at 3 and 7 months of age. However, at 5 months the latency to trial end of *Mecp2*+/- mice on the supplemented diet is significantly higher than *Mecp2*+/- mice on control diet, indicating amelioration of motor coordination. (B) No differences are seen in the total distanced travelled in the open field test at 3 and 5 months of age for any of the mice, over a 30-minute period. At 7 months of age, both *Mecp2*+/- animals on the control and vitamin D supplemented diet travelled less in the open field than *Mecp2*+/+ animals on the control diet. (C) At 3 months of age, *Mecp2*+/- on the control, and *Mecp2*+/- and *Mecp2*+/+ mice on vitamin D supplemented diet spend significantly less time in the center of the open field when compared to *Mecp2*+/+ animals on the control diet, indicating increased anxiety-like behavior. At 5 months of age, no differences are seen between any of the groups. However, at 7 months of age, only the *Mecp2*+/- mice on control diet have increased anxiety-like behavior when compared to their wild-type littermates. (D) At 3 months of age, both *Mecp2*+/- mice on the control and vitamin D supplemented diet spend more time in the open arm of the elevated plus maze when compared to *Mecp2*+/+ mice. At 5 months, however, only *Mecp2*+/- mice on the supplemented diet spend more time in the open arm of the maze when compared to *Mecp2*+/+ mice. At 7 months of age, no differences are seen between groups. *p < 0.05, **p < 0.01, ****p < 0.0001. NS = not significant. Two-way ANOVA with Tukey’s post hoc analysis. N = 12 – 17 per condition, genotype and age group, indicated by dots. Error bars: ± SEM.

Next, we measured anxiety-related behavior, a phenotype that has been previously reported in RTT patients (Barnes et al., 2015) and mouse models (Adachi et al., 2009; Meng et al., 2016; Philippe et al., 2018; Samaco et al., 2013; Vogel Ciernia et al., 2017). In the open-field, mice that spend less time in the center compared to the periphery are considered to display increased anxiety-like behavior (Lezak et al., 2017). We find that in early and mid-symptomatic stages, vitamin D supplementation does not improve anxiety-like behavior of *Mecp2*+/- mice (Fig. 4C), with *Mecp2*+/- mice spending less time in the center than wildtype littermates. However, at 7 months of age, *Mecp2*+/- mice on the supplemented diet do not display increased anxiety-like behavior when compared to wildtype, suggesting an improvement in anxiety-like behavior with vitamin D supplementation.

We also employed the elevated plus maze to assesses anxiety-like behavior. Animals that spend an increased amount of time in the closed arm of the elevated maze relative to the open arm are thought to have higher levels of anxiety-related behavior (Lezak et al., 2017). However, studies have found that *Mecp2* deficient mice spend more time in the open arm of the maze, indicating less anxiety-like behavior (Meng et al., 2016; Ribeiro and Macdonald, 2020; Samaco et al., 2013; Ure et al., 2016; Vogel Ciernia et al., 2017), in opposition to their behavior in the open-field.

Recently, a study found that clipping the whiskers of *Mecp2* mutant animals eliminates elevated plus maze deficits, suggesting that this test might assess sensory hypersensitivity of RTT mice and not anxiety-related behavior (Flores Gutiérrez et al., 2020). We also find that *Mecp2*+/- mice spend more time in the open arm of the elevated plus maze than *Mecp2+/+* mice, indicating decreased anxiety-like behavior (or increased sensory hypersensitivity) in this paradigm; vitamin D supplementation does not alter this phenotype, however (Fig. 4D).

Alterations in social interaction of *Mecp2* mutant mice are dependent on strain and background (Ribeiro and Macdonald, 2020; Vogel Ciernia et al., 2017). To examine this behavior in our mice, we used the three-chambered social approach test in which mice are considered to have impaired social behavior when they spend more time with a novel object than an age- and sex-matched novel mouse (Yang et al., 2011). *Mecp2+/-* female mice on control or supplemented diet do not exhibit altered sociability when compared to *Mecp2+/+* mice (Supplemental Fig. 1). Taken together, these data indicated that dietary vitamin D supplementation improves motor coordination deficits and increased anxiety-like behavior in *Mecp2*+/- mice.

### Vitamin D deficient diet does not significantly exacerbate behavioral phenotypes

We next hypothesized that treating *Mecp2*+/- mice with a diet deficient in vitamin D (0.1 IU/g) would exacerbate their behavioral phenotypes. To test this hypothesis, we followed the same experimental design used in the supplementation study, placing *Mecp2*+/- mice and wildtype *Mecp2*+/+ littermates on 0.1 IU/g vitamin D diet at weaning (1 month of age), and performing the behavioral tests at 3, 5, and 7 months of age. At all time-points on the accelerating rotarod, *Mecp2*+/- females on both control and deficient vitamin D diets have a shorter latency to trial end compared to their *Mecp2*+/+ littermates, with no significant difference between those on deficient versus control diet (Fig. 5A). In the open field, we observe no difference among groups in the distance travelled at 3 and 5 months of age (Fig. 5B). At 7 months of age, there is a significant reduction in exploratory motor behavior and average speed of *Mecp2*+/- on both control and deficient diet, when compared to *Mecp2*+/+ on control diet (Fig. 5B). Interestingly, there is a trend to decreased distance traveled for *Mecp2*+/+ on the deficient diet, when compared to those on control diet. Because of this impact on wildtype mice, *Mecp2*+/- mice on the deficient diet do not show significant differences from their wildtype littermates on deficient diet.

**Figure 5.**
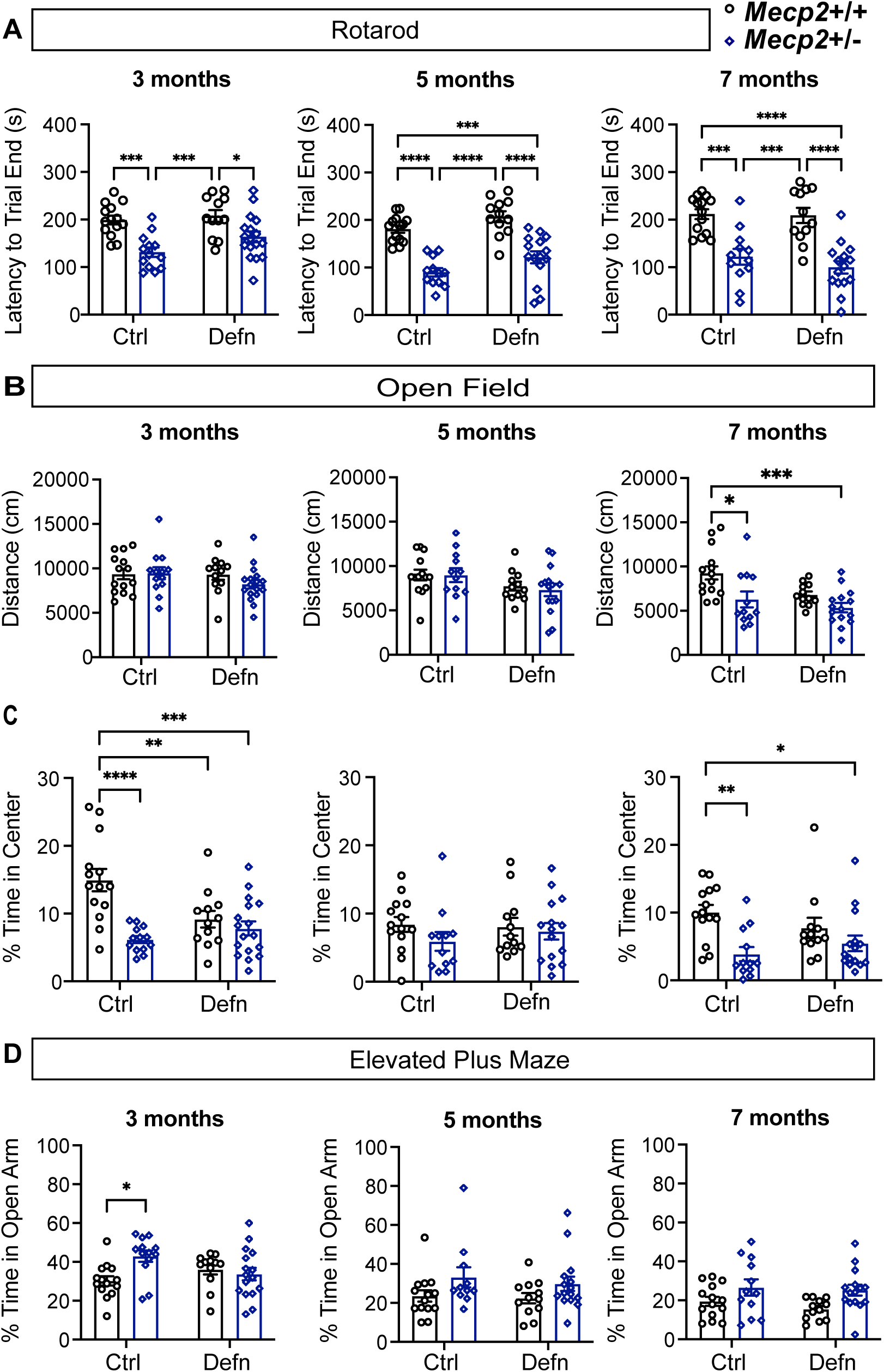
*Mecp2*+/- mice treated with vitamin D deficient and control diet display similar behavioral deficits. (A) In the rotarod test, *Mecp2*+/- mice on the control (Ctrl) and vitamin D deficient diet (Defn) have lower latency to trial end when compared to their wild-type (*Mecp2*+/+) littermates. At 5 and 7 months of age, *Mecp2*+/- mice on the control and deficient diet have a shorter latency to trial end than *Mecp2*+/+ animals. (B) In the open filed test, both *Mecp2*+/- mice on the control and deficient diet demonstrate impaired motor activity at 7 months of age, while no deficit is observed at 3 and 5 months of age. (C) At 3 months of age, *Mecp2*+/- on the control, *Mecp2*+/- and *Mecp2*+/+ mice on vitamin D deficient diet display increased anxiety-like behavior when compared to *Mecp2*+/+ animals on the control diet, defined as reduced time in the center of the open field. At 5 months of age, no differences are seen between any of the groups. Importantly, at 7 months of age *Mecp2*+/- on both diets display increased anxiety-like behavior. (D) In the elevated plus maze, only the *Mecp2*+/- on the control diet display higher anxiety-like behavior than their wild-type littermates at 3 months of age. At 5 and 7 months of age, no differences are observed between animals treated with the control or vitamin D deficient diet. *p < 0.05, **p < 0.01, ***p < 0.001, ****p < 0.0001. Two-way ANOVA with Tukey’s post hoc analysis. N = 11 – 15 per condition and genotype, indicated by dots. Error bars: ±SEM.

A similar trend is observed in the percent time in center of the open-field; *Mecp2*+/- mice on deficient diet show a significant reduction in time in center compared to wildtype on control chow, but no significant differences from wildtype on deficient diet. It is important to note that at 7 months of age, wildtype mice on deficient diet show increased anxiety-like behavior compared to wildtype on control chow (Fig. 5C). *Mecp2*+/- mice on deficient chow do not behave significantly different than *Mecp2*+/- on control chow in the elevated plus maze (Fig. 5D) or 3 chamber sociability (Supplemental Fig. 1). Thus, our data show that exposure to a vitamin D deficient diet does not lead to an exacerbation of *Mecp2*+/- behavioral phenotypes; however, it does lead to some disruptions in wildtype behavior.

### Improved behavior in *Mecp2+/-* mice correlates with sufficient circulating vitamin D (25(OH)D) serum levels

We analyzed vitamin D serum concentrations at the end of the experiment (7 months) to determine whether the dietary modification of vitamin D resulted in changes in circulating 25(OH)D serum levels. We measured serum 25(OH)D (calcidiol) because it has been shown to best correlate with levels of the activated form of vitamin D (1,25(OH)D; calcitriol) in the brain (Spach and Hayes, 2005). We found that the deficient diet led to a significant reduction in serum 25(OH)D levels, in both wildtype and *Mecp2*+/-. This result is consistent with previous studies (Belenchia et al., 2017; Comer et al., 1993; Fleet et al., 2008; Kasatkina et al., 2020; Rowling et al., 2007), and it is likely due to a reduction in the availability of 25(OH)D precursor. The supplemented diet, on the other hand, did not significantly alter 25(OH)D serum levels of *Mecp2+/-* or *Mecp2+/+* mice at 7 months of age; however, we observed high variability in 25(OH)D levels on both the control and supplemented diets (Fig. 6A). To investigate whether circulating levels of vitamin D correlate with behavioral phenotypes, which also display variability, we analyzed our data based on the 25(OH)D serum concentration, regardless of the diet they were treated with. We considered deficient levels to be below 20 ng/ml, insufficient levels to be between 20 and 29 ng/ml, and sufficient levels to be above 29 ng/ml (Holick et al., 2011; Mallya et al., 2016).

**Figure 6.**
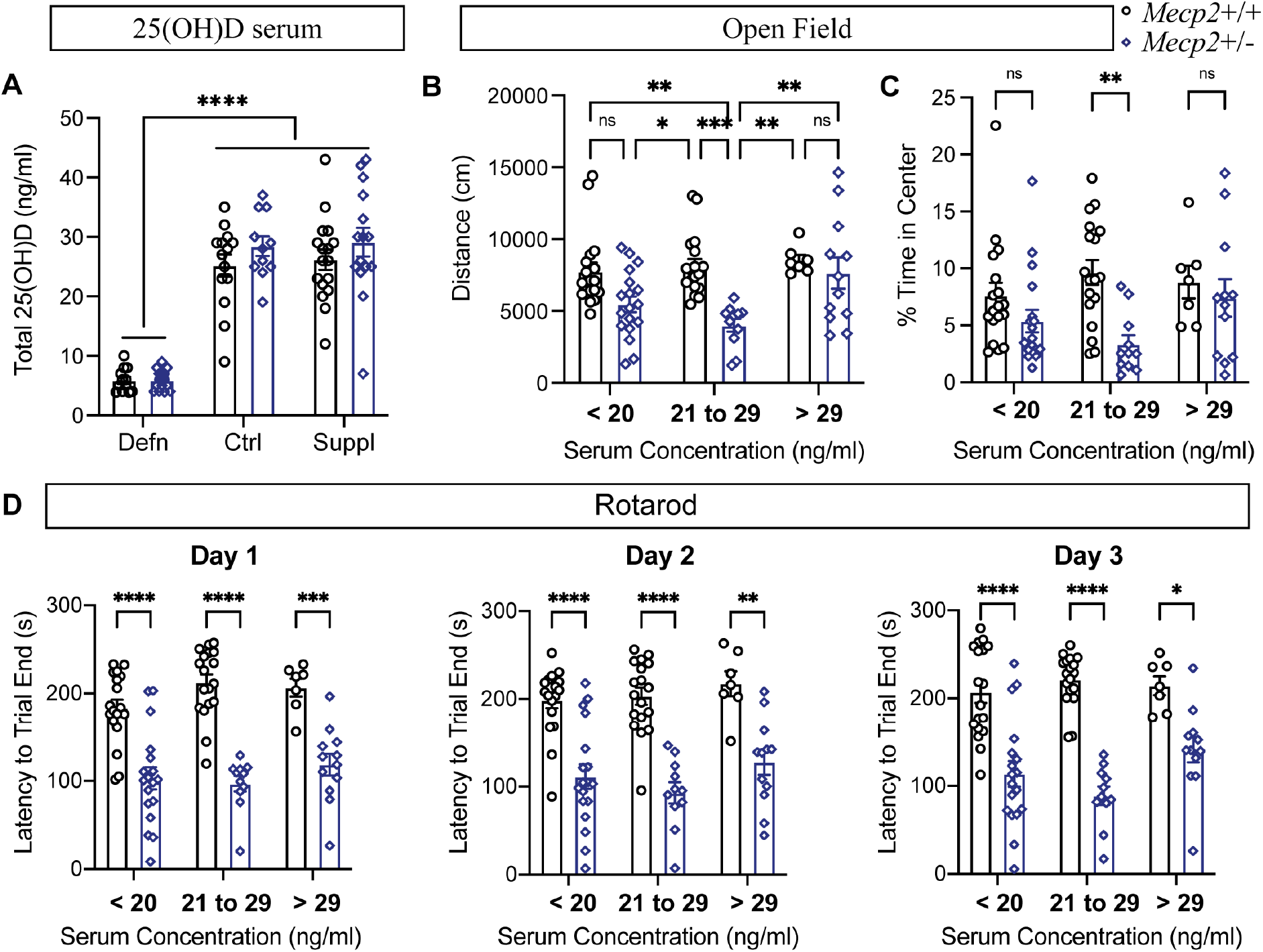
Sufficient serum vitamin D (25(OH)D) rescues motor activity and anxiety-like behavior of *Mecp2*+/- mice. (A) Total serum 25(OH)D (calcidiol) is not increased at 7 months of age in mice fed a vitamin D supplemented diet (Suppl), compared to those on the control diet (Ctrl), regardless of genotype. However, vitamin D deficient diet (Defn) significantly reduces 25(OH)D serum of mice, independent of genotype. N = 11 – 17 per condition and genotype, indicated by dots. (B-D) For these analyses, animals were grouped based on serum 25(OH)D concentration, and not dietary treatment. Serum 25(OH)D lower than 20 ng/ml is considered deficient, while 21 to 29 ng/ml is insufficient, and greater than 29 ng/ml is considered sufficient 25(OH)D vitamin D. (B) *Mecp2*+/- mice with insufficient serum 25(OH)D cover significantly less distance in the open field than wildtype. *Mecp2*+/- mice with sufficient serum 25(OH)D, however, cover significantly more distance in the open field than *Mecp2*+/- with insufficient levels, indicating a rescue in exploratory motor activity. (C) *Mecp2*+/- mice with insufficient serum 25(OH)D concentration display an increase in anxiety-like behavior when compared to their wild-type littermates, while *Mecp2*+/- mice with either deficient and sufficient levels do not display a significant difference in the percent time in the center of the open field. (D) *Mecp2*+/- mice with deficient, insufficient, and sufficient serum 25(OH)D display shorter latency to trial end when compared to *Mecp2*+/+ mice. No differences are seen between *Mecp2*+/+ in all three groups. *p < 0.05, **p < 0.01, ***p < 0.001, ****p < 0.0001. NS = not significant. Two-way ANOVA with Tukey’s post hoc analysis. B, C: N = 7 – 24 per condition and genotype, indicated by dots. Error bars: ± SEM.

We find that *Mecp2*+/- mice with insufficient levels of serum 25(OH)D travel significantly less in the open-field arena than *Mecp2+/+* mice in any group (Fig. 6B); however, *Mecp2*+/- mice with sufficient levels cover significantly more distance than those with insufficient levels and they are not significantly different from wildtype. The majority of mice with deficient vitamin D serum levels are those that were on the vitamin D deficient diet. In keeping with the results observed for the vitamin D deficient diet, *Mecp2*+/- with deficient serum levels do not display a significant reduction in distance traveled compared to deficient *Mecp2*+/+, although they do show a reduction relative to *Mecp2*+/+ with insufficient or sufficient levels. *Mecp2+/+* mice with insufficient 25(OH)D serum concentration are not impacted and display similar exploratory behavior to *Mecp2+/+* mice with sufficient levels, suggesting that wildtype mice are not as sensitive to insufficient levels of vitamin D as *Mecp2*+/- mice are.

In addition, *Mecp2+/-* female mice with sufficient 25(OH)D serum concentrations do not display increased anxiety-like behavior in the open field test, when compared to *Mecp2+/+* mice, while *Mecp2*+/- mice with insufficient 25(OH)D serum levels exhibit increased anxiety-like behavior (Fig. 6C). However, lower levels of serum 25(OH)D do not significantly alter behavior in the elevated plus maze (Supplemental Fig. 2) or social approach (Supplemental Fig. 2). Although serum 25(OH)D does not significantly alter the reduced motor coordination observed in *Mecp2*+/- mice on the accelerating rotarod (Fig. 6D), there is a trend toward less severe deficits in *Mecp2*+/- mice with sufficient levels, when compared to wildtype littermates. Thus, our results suggest that low levels of serum 25(OH)D in *Mecp2+/-* mice could contribute to exploratory motor and anxiety-like behavioral deficits.

### Vitamin D homeostasis is disrupted in multiple tissues of *Mecp2*+/- mice

In view of the similar effect that vitamin D supplementation and deficiency exerted on behavior of *Mecp2*+/- mice, as well as the partial overlap in rescue of DEGs, we investigated the hypothesis that vitamin D homeostasis might be altered in *Mecp2*+/- mice, leading to the activation of compensatory mechanisms. Vitamin D metabolism is complex and involves multiple genes and tissues (Fig. 7A). Vitamin D3 and D2 are processed into calcidiol (25(OH)D) in the liver by the cytochrome oxidases CYP27A1 and CYP2R1. Calcidiol is then transported to the kidney and processed into calcitriol (1,25(OH)2D3), the activated form of vitamin D, by the enzyme CYP27B1 (Zhu et al., 2013). Finally, the enzyme CYP24A1 is responsible for the catabolism of calcitriol (Jeon and Shin, 2018; Jones et al., 2014; Schuster, 2011). There is a feedback mechanism in place that adjusts the levels of these enzymes in response to vitamin D, maintaining homeostasis. For example, higher levels of serum 25(OH)D corresponds with lower levels of *CYP27B1* and upregulation of *CYP24A1* (Fleet et al., 2008). The cytochrome oxidase enzymes are also expressed in the brain (Gezen-Ak et al., 2013; Landel et al., 2018) and both forms of vitamin D, calcidiol and calcitriol, are able to cross the blood brain barrier (Pardridge et al., 1985), suggesting that the brain also synthesizes vitamin D *in situ* (Spach and Hayes, 2005).

**Figure 7.**
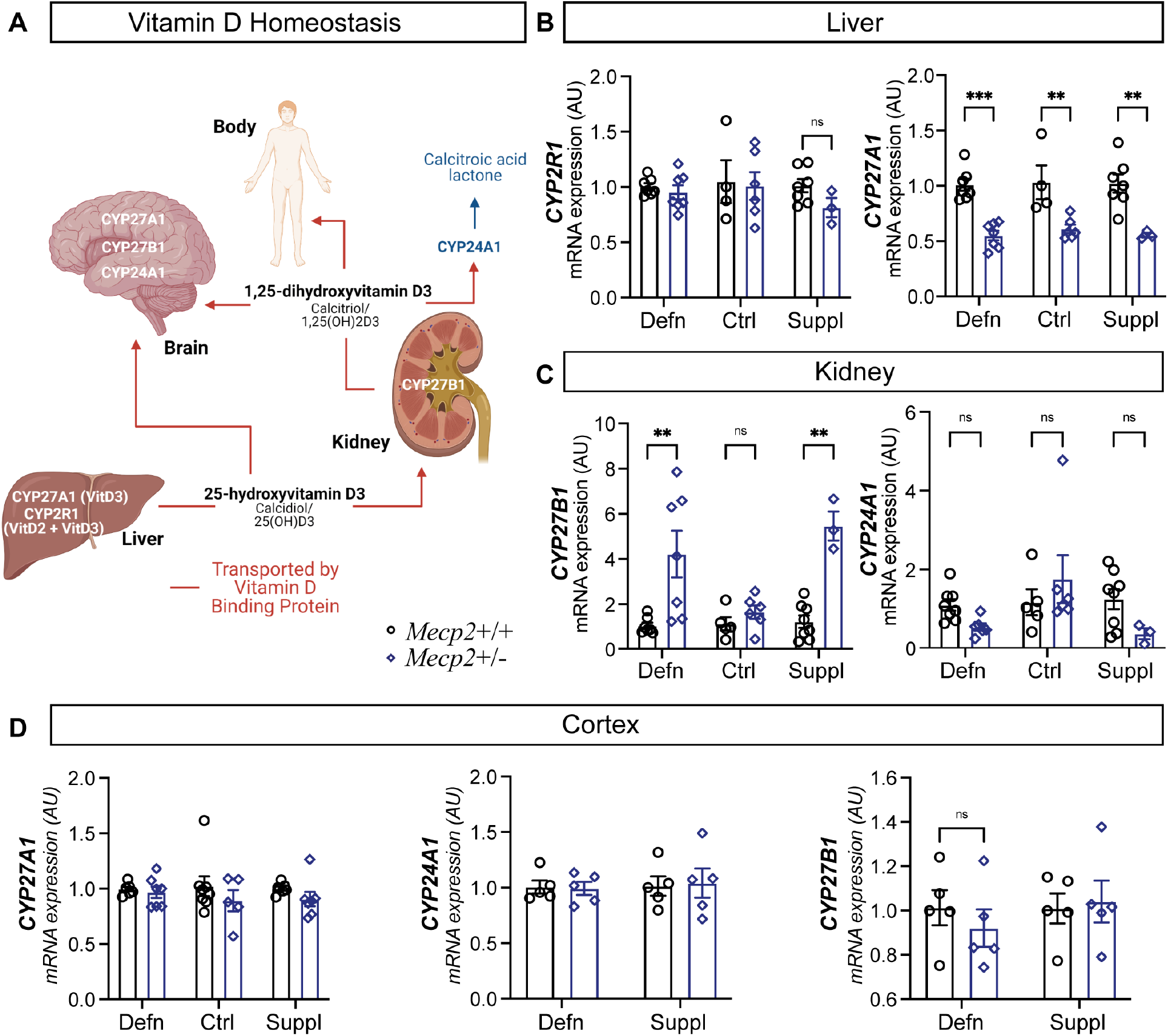
*Mecp2*+/- mice have disruptions in the vitamin D synthesis pathway and display disruptions in vitamin D homeostasis with dietary modulation. (A) Illustration of vitamin D biosynthesis process. In the liver, the enzymes CYP27A1 and CYP2R1 process vitamin D into 25(OH)D (calcidiol), which is then transported to the kidneys. The enzyme CYP27B1 hydrolyzes 25(OH)D into 1,25(OH)2 (calcitriol), which is transported to various organs or metabolized by the enzyme CYP24A1. The brain expresses the enzymes involved in the vitamin D metabolism, indicating synthesis *in situ*. (B) *CYP2R1* expression is unaltered with vitamin D dietary deficiency and supplementation. *Mecp2*+/- on all three diets display reduced expression of hepatic *CYP27A1*. (C) The expression of renal *CYP27B1* is elevated in *Mecp2*+/- mice on vitamin D supplemented and deficient diets, relative to *Mecp2*+/+, while no difference is seen in the kidney of *Mecp2*+/- animals on the control diet. Conversely, *CYP24A1* expression has a trend towards downregulation in the kidney of *Mecp2*+/- mice on both vitamin D deficient and supplemented diets, relative to *Mecp2*+/+. (D) No significant differences are observed in cortical gene expression of *CYP27A1*, *CYP24A1* and *CYP27B1* between *Mecp2*+/- and *Mecp2*+/+. **p < 0.01, ***p < 0.001. Two-way ANOVA with Tukey’s post hoc analysis. N = 3 – 8 per condition and genotype, indicated by dots. Error bars: ±SEM.

To investigate whether vitamin D homeostasis is disrupted in *Mecp2*+/- mice, we analyzed the expression of the genes involved in vitamin D metabolism. In the liver, there is no alteration in the expression of *CYP2R1*; however, *Mecp2+/-* females display reduced levels of *CYP27A1* expression compared to wildtype, regardless of vitamin D diet (Fig. 7B), suggesting disruptions in the synthesis of calcidiol. Strikingly, *Mecp2+/-* females on both the supplemented and deficient diets display elevated levels of *CYP27B1* and a trend towards lower *CYP24A1* expression in the kidney (Fig. 3.7C), while *Mecp2*+/- on the control diet do not have significant differences from wildtype. It is thus interesting to speculate that modulating vitamin D in the diet, in either direction, leads to an up-regulation in the synthesis and decrease in the catabolism of calcitriol in the kidney, which could compensate for the reduced production of calcidiol in the liver (*CYP27A1*), and that this could contribute to the similar behavioral phenotypes observed with deficient and supplemented diets. Interestingly, there is no change in the expression of these genes in the cortex of *Mecp2*+/- mice, however (Fig. 7D), indicating that the alterations in vitamin D homeostasis might be specific to peripheral tissues.

## Discussion

In this study, we investigated whether vitamin D restores homeostasis to dysregulated gene expression in the cortex of *Mecp2*+/- mice, and whether vitamin D impacts behavioral phenotypes in these RTT model mice. We found that dietary supplementation and deficiency of vitamin D have a broad impact on the transcriptome of the cortex, regardless of genotype; strikingly, we identified more than 200 genes whose expression is dysregulated in *Mecp2*+/- cortex under control conditions but rescued by dietary vitamin D supplementation. We further found that *Mecp2*+/- mice exhibit a disruption in the homeostasis of vitamin D metabolism in the periphery, and that multiple behaviors correlate with vitamin D serum levels, including motor activity and anxiety-like behavior in the open-field.

Vitamin D supplementation normalizes the expression of 283 dysregulated genes in the *Mecp2*+/- cortex, which we hypothesize underpin the rescue in neuronal morphology and behavior observed with vitamin D. Many of these rescued DEGs play important roles in neuronal morphology and brain function, including *CYP46A1*, the major contributor to cholesterol catabolism in the brain (Lund et al., 1999). Cholesterol is essential for proper brain function and it is synthesized *in situ*, as it cannot cross the blood-brain-barrier (Turley et al., 1998, 1996). Cholesterol metabolism is significantly altered in a number of conditions affecting the CNS, including RTT (reviewed in Martín et al., 2014). Downregulation of *CYP46A1* is observed in severely symptomatic male *Mecp2*-null mice, while at early symptomatic stages, its expression is increased (Buchovecky et al., 2014). We find that *CYP46A1* is downregulated in the cortex of *Mecp2*+/- mice on the control chow at 7 months, relative to wildtype, while vitamin D supplementation normalizes *CYP46A1* expression. Importantly, vitamin D deficiency exacerbates the reduced expression of *CYP46A1* in *Mecp2*+/- cortex. This is particularly compelling finding as dysregulation of *CYP46A1* plays an important role in other neurological disorders, and it is currently being studied as a target for therapeutics in Huntington’s (Boussicault et al., 2016; Kacher et al., 2019) and Alzheimer’s diseases (Chen et al., 2020; Djelti et al., 2015; Kölsch et al., 2009).

Dietary vitamin D deficiency leads to additional transcriptome differences between *Mecp2*+/+ and *Mecp2*+/- cortex, including exacerbating the dysregulation of 23 of the DEGs that are rescued by supplementation. According to GO analysis, these 23 genes are particularly involved in the regulation of neurotransmitter receptor activity, such as histamine receptors (HRH3) and GABA receptors (GABBR1). Unexpectedly, 54 of the DEGs rescued with vitamin D supplementation are also normalized with its deficiency. A few of these genes have been associated with neurodevelopmental disorders; for example, copy number variants in KANK1 are a risk factor for ASD (Vanzo et al., 2019), and ANK2 is a high-confidence ASD gene (Iossifov et al., 2014; Yang et al., 2019). These data suggest that alterations in vitamin D homeostasis, whether through supplementation or restriction, normalizes the expression of a select number of clinically relevant genes in the *Mecp2*+/- cortex. It is important to note, however, that none of the DEGs whose expression is rescued on the deficient diet contain the conserved VDR response element motif, suggesting that their transcription is not directly regulated by vitamin D and the VDR.

Vitamin D does not act exclusively in a genomic (direct transcriptional) capacity, however; PDIA3 is a vitamin D receptor that is localized to the cell membrane and activates downstream signaling cascades in response to vitamin D (Boyan et al., 2012; Chen et al., 2010; J. Chen et al., 2013). We have shown previously that *Pdia3* expression is unaltered in the cortices of male and female *Mecp2* mutant mice at 2 and 5 months of age, respectively (Ribeiro et al., 2020). Our RNA-sequencing data confirm this result in *Mecp2*+/- on control chow at 7 months of age. Nonetheless, with vitamin D supplementation, *Pdia3* expression increases in *Mecp2*+/- cortex, when compared to both wild-type and *Mecp2*+/- mice on the control chow. Interestingly, vitamin D deficiency results in significant up-regulation of *Pdia3* in both *Mecp2*+/- and *Mecp2*+/+ cortex. This apparent compensatory up-regulation of *Pdia3* in response to both supplementation and deficiency of vitamin D could explain the overlap in rescue of DEGs, and perhaps contribute to the lack of exacerbation of behavioral phenotypes with vitamin D deficiency.

Serum measurements of 25(OH)D collected at 7 months confirmed the efficacy of the vitamin D deficient diet (0.1 IU/g vitamin D), with all mice showing vitamin D deficiency, defined by serum 25(OH)D levels below 20 mg/ml. Mice on the vitamin D supplemented diet (10 IU/g), on the other hand, showed greater variability in serum 25(OH)D levels. Overall, there was not a significant increase in serum 25(OH)D levels for mice (of either genotype) on the supplemented diet compared to the control (1 IU/g) diet. We previously identified an increase in serum 25(OH)D levels with 10 IU/g vitamin D supplementation at 5 months of age, independent of genotype (Ribeiro et al., 2020). This could explain the rescue of motor coordination behavior observed on the rotarod at 5 months but not 7 months. One possible explanation for the lack of significant increase in serum vitamin D at 7 months compared to 5 months is an increase in body weight during this time. Studies have shown that obesity can diminish the effects of vitamin D supplementation (Castaneda et al., 2012; Gallagher et al., 2013; Lee et al., 2009; Wortsman et al., 2000). The mechanism responsible for this is still poorly understood, but it might involve volumetric dilution of vitamin D in a large body mass (Drincic et al., 2012) or fat sequestration (Wortsman et al., 2000). Our data show that *Mecp2*+/- mice are significantly heavier at 7 months of age than at 5 months (Supplemental Fig. 3); therefore, it is possible that *Mecp2*+/- mice might need a higher dose of vitamin D later in life in order to enhance 25(OH)D serum concentration to compensate for the increase in body weight. However, we do not observe an overall correlation between 25(OH)D status and body weight at 7 months of age, consistent with previous findings (Seldeen et al., 2017).

Serum vitamin D status does appear to correlate with behavior in the open-field, however. *Mecp2*+/- mice with sufficient levels of vitamin D (> 29 ng/ml) travel the same distance and have the same overall velocity as wildtype mice. Additionally, they spend the same amount of time in the center of the open-field. *Mecp2*+/- mice with insufficient levels (< 29 ng/ml), on the other hand, travel significantly less in the open-field, have a reduced velocity, and spend significantly less time in the center of the open-field than wildtype or mice with sufficient vitamin D. We hypothesized that we would observe an exacerbation of behavioral phenotypes in *Mecp2*+/- mice with dietary vitamin D deficiency, but surprisingly found that those on the deficient diet performed better in some tests than those on the control diet.

One possible explanation for the influence of low 25(OH)D concentration on behavior lies within vitamin D homeostasis. Expression of *CYP27A1*, a rate-limiting factor in the production of 25(OH)D, is reduced in the liver of *Mecp2*+/- mice on all chows. The reduced expression of *CYP27A1* in the liver in *Mecp2*+/- mice on supplemented chow could explain why renal *CYP27B1* expression is upregulated in response to vitamin D supplementation; greater concentration of 25(OH)D precursor could lead to increased expression of the gene required for the production of the active metabolite, in an effort to normalize vitamin D metabolism. There is also a trend toward reduced *CYP24A1* in the liver, the enzyme responsible for catabolizing 1,25(OH)_2_D, which could also help normalize activated vitamin D levels. However, since *CYP27A1* is a limiting factor, vitamin D homeostasis remains disrupted in the *Mecp2*+/- mice. Because *Mecp2*+/- on the vitamin D deficient diet also display reduced *CYP27A1* expression in the liver, increased *CYP27B1* and a trend towards decrease *CYP24A1* expression in the kidney, these mice might also benefit from a compensatory mechanism leading to increased activated circulating vitamin D relative to those on control diet. These data highlight the complexity of vitamin D, and the important cross-talk between the periphery and the brain that contributes to RTT phenotypes.

RTT is a highly complex disorder, caused by myriad transcriptome and proteome disruptions that occur downstream of mutations in MeCP2. Thus, treating RTT will likely require a combinatorial approach, restoring homeostasis to multiple disrupted cellular pathways that additively contribute to RTT phenotypes. Vitamin D has the ability to modify multiple of these disrupted cellular pathways in parallel. Here, we show that dietary vitamin D has a profound impact on the cortical transcriptome, and that it is associated with changes in behavioral phenotypes in *Mecp2*+/- mice. Although further studies on cholesterol and vitamin D metabolism in the brain of *Mecp2* mutant mice are warranted to fully understand the broad mechanism of action of vitamin D dietary modulation, it has the potential to provide valuable benefit to the quality of life of RTT patients.

## Declarations of competing interest

none

## Acknowledgements

The authors thank Dr. Yasir Ahmed-Braimah (Syracuse University) for assistance with RNA-seq experiments and data analysis, Leanne Kelley for assistance with data analysis, and Dr. Sarah Hall (Syracuse University) for critical feedback on the manuscript. This work was supported by the National Institutes of Health [Grant number 1R01NS106285], and the International Rett Syndrome Foundation [Grant number 3064] awarded to JLM.

## Author Contributions

MCR and JLM designed the experiments, MCR performed experiments, MCR and JLM analyzed data and wrote and edited the manuscript.

## Notes

### Competing Interest Statement

The authors have declared no competing interest.

