## Supplemental for "Vitamin D modulates cortical transcriptome and behavioral phenotypes in an *Mecp2* heterozygous Rett syndrome mouse model"

Department of Biology

360 Life Sciences Complex

107 College Place

Syracuse University

Syracuse, NY 13244

**Supplemental Figures: 3**

**Supplemental Tables: 4**

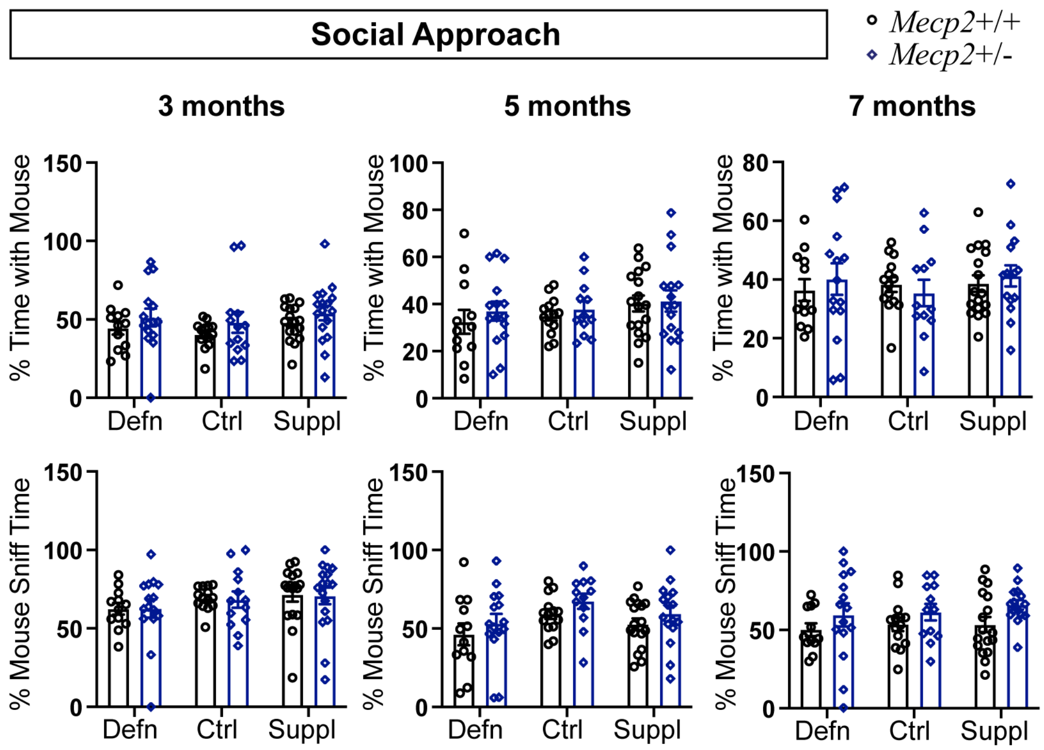


**Supplemental Figure 1. Vitamin D modulation does not alter sociability of *Mecp2*+/- mice.** We observe no difference in social behavior in any of the studied groups at 3, 5 and 7 months of age, as indicated by percent time spent investigating a novel mouse and percent time sniffing the novel mouse. Two-way ANOVA with Tukey's post hoc analysis. N = 11 – 17 per condition, genotype, and age group, indicated by dots. Error bars: ± SEM.

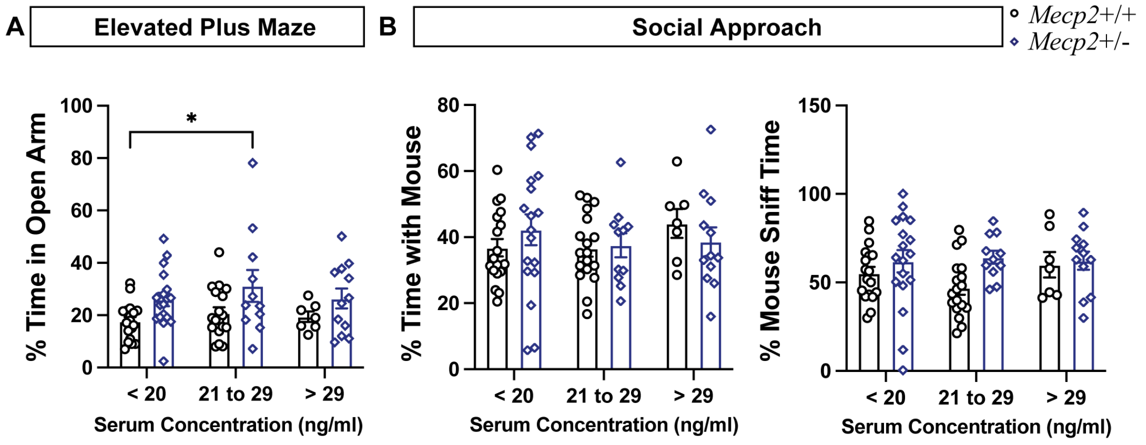


**Supplemental Figure 2. Serum 25(OH)D concentration does not alter behavioral phenotypes in the elevated plus maze or social approach tests.**

(A) Only *Mecp2*+/- mice with sufficient serum 25(OH)D concentration display increased percent time spent in the open arm when compared to wild-type mice with deficient serum 25(OH)D. (B) No differences were observed in the social approach test for any of the groups, as indicated by percent time spent investigating a novel mouse and percent time sniffing the novel mouse. $^*p < 0.05$. Two-way ANOVA with Tukey's post hoc analysis. N = 11 – 15 per condition and genotype, indicated by dots. Error bars: ± SEM.

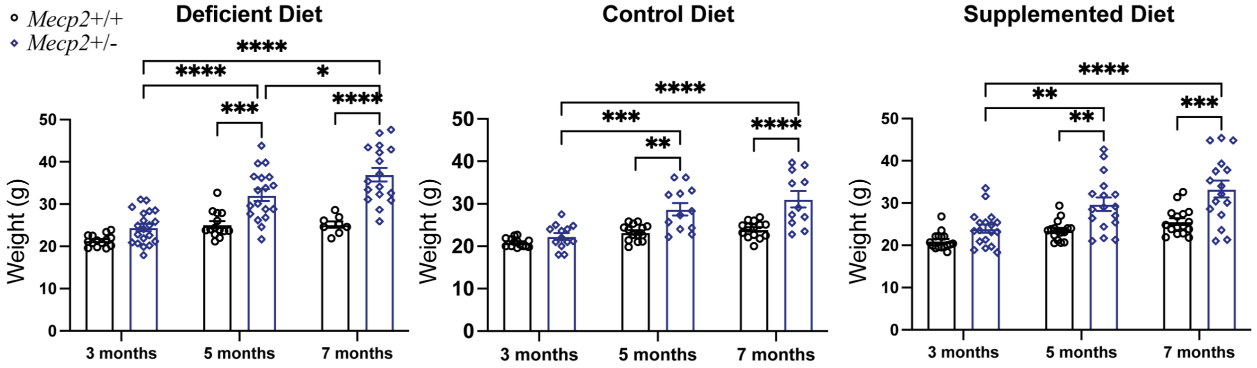


**Supplemental Figure 3. *Mecp2*+/- mice have increased weight in late symptomatic stage, regardless of vitamin D.**

At 3 months of age *Mecp2*+/- mice do not weight significantly more than *Mecp2*+/+ littermates on any of the diets. At 5 and 7 months of age, however, *Mecp2*+/- on all diets exhibit increased weight, relative to *Mecp2*+/+. $^*p < 0.05$, $^{**}p < 0.01$, $^{***}p < 0.001$, $^{****}p < 0.0001$. Two-way ANOVA with Tukey's post hoc analysis. N = 11 – 15 per condition and genotype, indicated by dots. Error bar: ±SEM.

**Table 1.** DEGs in *Mecp2*+/- cortex, relative to *Mecp2*+/+, that are normalized with vitamin D supplementation and differentially expressed in the cortex of *Mecp2*+/- on supplemented vs control diet.

| | **Ctrl *Mecp2*+/- vs *Mecp2*+/+** | | | ***Mecp2*+/- Suppl vs Ctrl** | | |
|---|---|---|---|---|---|---|
| **Gene ID** | **P-value** | **FDR** | **Fold change** | **P-value** | **FDR** | **Fold change** |
| 0610012G03Rik | 0.0034 | 0.0817 | -1.1559 | 0.0001 | 0.0004 | 1.2620 |
| 2010204K13Rik | 0.0020 | 0.0681 | -1.1899 | 0.0471 | 0.0941 | 1.1062 |
| 4933432I09Rik | 0.0055 | 0.0998 | -2.7857 | 0.0159 | 0.0391 | 2.2857 |
| A2ml1 | 0.0031 | 0.0794 | -1.2304 | 0.0004 | 0.0020 | 1.3045 |
| A330009N23Rik | 0.0026 | 0.0734 | -1.2352 | 0.0001 | 0.0004 | 1.3888 |
| A730085K08Rik | 0.0002 | 0.0309 | -2.1800 | 0.0003 | 0.0016 | 2.0667 |
| Abca2 | 0.0008 | 0.0495 | -1.2735 | 0.0001 | 0.0008 | 1.3404 |
| Abi3 | 0.0043 | 0.0900 | -1.2380 | 0.0047 | 0.0144 | 1.2334 |
| Abitram | 0.0040 | 0.0871 | -1.2738 | 0.0006 | 0.0027 | 1.3570 |
| Add1 | 0.0009 | 0.0518 | -1.1679 | 0.0002 | 0.0013 | 1.1963 |
| Aff1 | 0.0017 | 0.0637 | 1.3975 | 0.0030 | 0.0100 | -1.3573 |
| Agpat4 | 0.0028 | 0.0755 | -1.1728 | 0.0013 | 0.0052 | 1.1902 |
| Ajm1 | 0.0003 | 0.0353 | -1.2287 | 0.0000 | 0.0001 | 1.3468 |
| Anapc13 | 0.0039 | 0.0865 | -1.1887 | 0.0022 | 0.0079 | 1.2076 |
| Ank2 | 0.0006 | 0.0459 | 1.2390 | 0.0004 | 0.0019 | -1.2556 |
| Ankrd13b | 0.0038 | 0.0855 | -1.3211 | 0.0026 | 0.0090 | 1.3422 |
| Ap1s1 | 0.0003 | 0.0353 | -1.1961 | 0.0002 | 0.0011 | 1.2068 |
| Apln | 0.0026 | 0.0734 | -1.2239 | 0.0375 | 0.0782 | 1.1378 |
| Aqr | 0.0024 | 0.0713 | 1.2618 | 0.0011 | 0.0045 | -1.2881 |
| Arfgef2 | 0.0043 | 0.0895 | 1.3171 | 0.0149 | 0.0372 | -1.2607 |
| Arhgap33 | 0.0014 | 0.0589 | -1.3896 | 0.0111 | 0.0294 | 1.2653 |
| Arsa | 0.0013 | 0.0585 | 1.5123 | 0.0011 | 0.0043 | -1.5236 |
| Atn1 | 0.0008 | 0.0495 | 2.3771 | 0.0011 | 0.0043 | -2.3376 |
| Atp6ap1l | 0.0054 | 0.0986 | -1.3218 | 0.0099 | 0.0266 | 1.2870 |
| AW209491 | 0.0040 | 0.0871 | 1.3951 | 0.0140 | 0.0355 | -1.3082 |
| Azi2 | 0.0041 | 0.0871 | 1.1646 | 0.0002 | 0.0011 | -1.2448 |
| B3galt2 | 0.0014 | 0.0599 | 1.2108 | 0.0150 | 0.0375 | -1.1456 |
| B4galnt1 | 0.0012 | 0.0548 | -1.2064 | 0.0319 | 0.0687 | 1.1183 |
| Bcan | 0.0002 | 0.0297 | -1.2240 | 0.0000 | 0.0000 | 1.3260 |
| Bcl7a | 0.0001 | 0.0274 | 1.4932 | 0.0077 | 0.0217 | -1.2732 |
| Btbd17 | 0.0001 | 0.0274 | -1.3179 | 0.0002 | 0.0012 | 1.2899 |
| C1ra | 0.0000 | 0.0186 | -1.7489 | 0.0008 | 0.0034 | 1.4978 |
| C2cd5 | 0.0000 | 0.0165 | 1.4859 | 0.0001 | 0.0006 | -1.3971 |
| Cacnb3 | 0.0011 | 0.0547 | -1.2627 | 0.0018 | 0.0066 | 1.2481 |

| | | | | | | |
|---|---|---|---|---|---|---|
| Cacnb4 | 0.0000 | 0.0186 | 1.4548 | 0.0000 | 0.0000 | -1.6435 |
| Caly | 0.0000 | 0.0188 | -1.2074 | 0.0000 | 0.0003 | 1.2107 |
| Camk1d | 0.0007 | 0.0462 | 1.2202 | 0.0010 | 0.0040 | -1.2078 |
| Camkv | 0.0025 | 0.0713 | -1.1997 | 0.0012 | 0.0047 | 1.2243 |
| Car11 | 0.0009 | 0.0518 | -1.1934 | 0.0005 | 0.0023 | 1.2102 |
| Carmil3 | 0.0029 | 0.0778 | -1.3377 | 0.0132 | 0.0339 | 1.2691 |
| Cbr1 | 0.0029 | 0.0767 | -1.1298 | 0.0156 | 0.0386 | 1.0982 |
| Cbs | 0.0010 | 0.0534 | -1.3784 | 0.0000 | 0.0002 | 1.6022 |
| Ccdc116 | 0.0054 | 0.0986 | -2.1786 | 0.0361 | 0.0759 | 1.7143 |
| Ccdc3 | 0.0022 | 0.0692 | -1.3728 | 0.0485 | 0.0963 | 1.2156 |
| Ccnb2 | 0.0030 | 0.0781 | -2.3125 | 0.0003 | 0.0013 | 3.0000 |
| Cd34 | 0.0047 | 0.0933 | -1.2541 | 0.0022 | 0.0079 | 1.2884 |
| Cd47 | 0.0001 | 0.0222 | 1.5120 | 0.0012 | 0.0046 | -1.3562 |
| Cda | 0.0015 | 0.0618 | -2.1837 | 0.0015 | 0.0057 | 2.2041 |
| Cdk2ap2 | 0.0009 | 0.0505 | -1.2910 | 0.0002 | 0.0012 | 1.3477 |
| Cdk5 | 0.0021 | 0.0685 | -1.1565 | 0.0006 | 0.0026 | 1.1842 |
| Cebpg | 0.0034 | 0.0817 | 1.2351 | 0.0001 | 0.0006 | -1.3705 |
| Chid1 | 0.0002 | 0.0336 | -1.2587 | 0.0070 | 0.0201 | 1.1594 |
| Chl1 | 0.0039 | 0.0860 | -1.1598 | 0.0002 | 0.0013 | 1.2325 |
| Chmp1b | 0.0010 | 0.0527 | 1.2903 | 0.0103 | 0.0277 | -1.2008 |
| Chpf2 | 0.0030 | 0.0781 | -1.1798 | 0.0002 | 0.0013 | 1.2476 |
| Cic | 0.0043 | 0.0903 | -1.1875 | 0.0052 | 0.0156 | 1.1779 |
| Clic1 | 0.0027 | 0.0741 | -1.6460 | 0.0272 | 0.0605 | 1.3982 |
| Clptm1 | 0.0020 | 0.0681 | -1.1790 | 0.0117 | 0.0306 | 1.1359 |
| Comt | 0.0006 | 0.0452 | 1.2714 | 0.0136 | 0.0346 | -1.1693 |
| Cops9 | 0.0023 | 0.0704 | -1.1729 | 0.0386 | 0.0802 | 1.1046 |
| Cpeb2 | 0.0040 | 0.0871 | 1.3661 | 0.0001 | 0.0005 | -1.6177 |
| Cpne5 | 0.0002 | 0.0337 | -1.2708 | 0.0004 | 0.0020 | 1.2484 |
| Cpne6 | 0.0011 | 0.0542 | -1.2647 | 0.0030 | 0.0100 | 1.2392 |
| Cpsf1 | 0.0034 | 0.0824 | -1.1962 | 0.0002 | 0.0013 | 1.2743 |
| Creb5 | 0.0013 | 0.0582 | 1.6466 | 0.0035 | 0.0113 | -1.5711 |
| Crip2 | 0.0045 | 0.0919 | -1.2591 | 0.0478 | 0.0952 | 1.1604 |
| Cul9 | 0.0014 | 0.0595 | -1.3107 | 0.0000 | 0.0003 | 1.4929 |
| Cx3cr1 | 0.0000 | 0.0165 | -1.2899 | 0.0000 | 0.0000 | 1.3785 |
| Cyp46a1 | 0.0000 | 0.0165 | -1.2368 | 0.0000 | 0.0001 | 1.2407 |
| D430019H16Rik | 0.0033 | 0.0815 | -1.2226 | 0.0248 | 0.0560 | 1.1536 |
| Dbi | 0.0004 | 0.0406 | -1.2027 | 0.0001 | 0.0007 | 1.2417 |
| Dctpp1 | 0.0004 | 0.0405 | -1.2918 | 0.0041 | 0.0129 | 1.2149 |
| Ddx50 | 0.0019 | 0.0664 | 1.1851 | 0.0026 | 0.0090 | -1.1752 |
| Dhrs1 | 0.0031 | 0.0789 | -1.1754 | 0.0001 | 0.0006 | 1.2719 |

| | | | | | | |
|---|---|---|---|---|---|---|
| Dhx40 | 0.0026 | 0.0734 | -1.1963 | 0.0212 | 0.0494 | 1.1362 |
| Dipk1b | 0.0015 | 0.0618 | -1.1766 | 0.0024 | 0.0084 | 1.1668 |
| Dnlz | 0.0049 | 0.0941 | -1.1468 | 0.0206 | 0.0484 | 1.1131 |
| Egln2 | 0.0036 | 0.0836 | -1.1486 | 0.0017 | 0.0063 | 1.1666 |
| Eif2a | 0.0040 | 0.0871 | 1.2065 | 0.0002 | 0.0012 | -1.3091 |
| Ets1 | 0.0049 | 0.0941 | 1.4762 | 0.0002 | 0.0012 | -1.7500 |
| Fam155a | 0.0012 | 0.0563 | 1.1620 | 0.0002 | 0.0012 | -1.2022 |
| Fam171a2 | 0.0036 | 0.0832 | -1.1879 | 0.0001 | 0.0006 | 1.3009 |
| Fam20c | 0.0015 | 0.0618 | -1.2435 | 0.0001 | 0.0004 | 1.3729 |
| Fbn2 | 0.0043 | 0.0900 | -1.8750 | 0.0088 | 0.0241 | 1.7500 |
| Fbxo34 | 0.0006 | 0.0442 | 1.3352 | 0.0000 | 0.0003 | -1.4886 |
| Fcgr3 | 0.0048 | 0.0941 | -1.3060 | 0.0320 | 0.0689 | 1.2077 |
| Fcho2 | 0.0007 | 0.0469 | 1.2943 | 0.0004 | 0.0020 | -1.3140 |
| Fcrls | 0.0000 | 0.0165 | -1.3865 | 0.0003 | 0.0014 | 1.2860 |
| Fkbp1a | 0.0000 | 0.0165 | -1.2024 | 0.0000 | 0.0001 | 1.2309 |
| Fndc5 | 0.0013 | 0.0585 | -1.2299 | 0.0002 | 0.0011 | 1.2937 |
| Folh1 | 0.0047 | 0.0933 | 1.7647 | 0.0291 | 0.0637 | -1.5190 |
| Frmpd3 | 0.0003 | 0.0362 | 2.4385 | 0.0024 | 0.0085 | -2.0719 |
| Ftl1-ps1 | 0.0005 | 0.0406 | -1.3097 | 0.0084 | 0.0233 | 1.2003 |
| G3bp2 | 0.0007 | 0.0468 | 1.2003 | 0.0007 | 0.0031 | -1.1988 |
| Gabbr1 | 0.0037 | 0.0848 | -1.1223 | 0.0009 | 0.0039 | 1.1476 |
| Gabpb2 | 0.0005 | 0.0413 | 1.4481 | 0.0293 | 0.0641 | -1.2253 |
| Gabra4 | 0.0039 | 0.0865 | 1.1975 | 0.0006 | 0.0026 | -1.2678 |
| Gabrd | 0.0026 | 0.0734 | -1.1697 | 0.0001 | 0.0005 | 1.2636 |
| Gclc | 0.0041 | 0.0882 | 1.2387 | 0.0245 | 0.0555 | -1.1700 |
| Gdap1 | 0.0048 | 0.0941 | 1.1771 | 0.0239 | 0.0545 | -1.1364 |
| Gdf11 | 0.0018 | 0.0649 | -1.3043 | 0.0003 | 0.0016 | 1.3804 |
| Gli3 | 0.0023 | 0.0706 | 1.5858 | 0.0001 | 0.0007 | -1.8631 |
| Gm10076 | 0.0009 | 0.0505 | -1.1690 | 0.0343 | 0.0727 | 1.0913 |
| Gm10860 | 0.0014 | 0.0592 | -2.1765 | 0.0118 | 0.0307 | 1.8235 |
| Gm12543 | 0.0003 | 0.0353 | 1.6611 | 0.0031 | 0.0103 | -1.4802 |
| Gm14399 | 0.0047 | 0.0933 | -2.1176 | 0.0418 | 0.0854 | 1.6471 |
| Gm15337 | 0.0010 | 0.0525 | -1.6774 | 0.0043 | 0.0134 | 1.5376 |
| Gm16485 | 0.0042 | 0.0889 | 1.2671 | 0.0036 | 0.0115 | -1.2697 |
| Gm20712 | 0.0048 | 0.0933 | -1.2305 | 0.0007 | 0.0029 | 1.3018 |
| Gm42846 | 0.0020 | 0.0681 | -1.3237 | 0.0035 | 0.0114 | 1.3045 |
| Gm48283 | 0.0030 | 0.0785 | -1.2982 | 0.0004 | 0.0021 | 1.3925 |
| Gm50402 | 0.0010 | 0.0527 | -2.5000 | 0.0055 | 0.0165 | 2.1250 |
| Gm5576 | 0.0040 | 0.0871 | 2.5455 | 0.0015 | 0.0058 | -2.8000 |
| Gm9493 | 0.0043 | 0.0900 | 2.5283 | 0.0095 | 0.0258 | -2.1967 |

| | | | | | | |
|---|---|---|---|---|---|---|
| Gnas | 0.0016 | 0.0621 | 1.2507 | 0.0064 | 0.0185 | -1.2019 |
| Gpbp1l1 | 0.0038 | 0.0857 | 1.2073 | 0.0274 | 0.0607 | -1.1449 |
| Gpsm1 | 0.0028 | 0.0757 | -1.3461 | 0.0015 | 0.0057 | 1.3916 |
| Gpt | 0.0030 | 0.0781 | -1.3894 | 0.0132 | 0.0338 | 1.3135 |
| Grcc10 | 0.0005 | 0.0427 | -1.1605 | 0.0000 | 0.0002 | 1.2239 |
| Grina | 0.0010 | 0.0525 | -1.1810 | 0.0000 | 0.0002 | 1.2718 |
| Grk6 | 0.0016 | 0.0629 | -1.1921 | 0.0001 | 0.0007 | 1.2679 |
| Grm8 | 0.0021 | 0.0685 | 1.2604 | 0.0001 | 0.0008 | -1.3725 |
| Gstm2 | 0.0037 | 0.0840 | -1.5324 | 0.0231 | 0.0529 | 1.3813 |
| Gstp1 | 0.0054 | 0.0986 | -1.1233 | 0.0022 | 0.0079 | 1.1400 |
| Gucy2e | 0.0038 | 0.0857 | -1.6019 | 0.0134 | 0.0342 | 1.4563 |
| H2-DMa | 0.0025 | 0.0725 | -1.4063 | 0.0002 | 0.0012 | 1.5596 |
| H2aj | 0.0018 | 0.0659 | -1.2572 | 0.0223 | 0.0515 | 1.1651 |
| Haus2 | 0.0043 | 0.0900 | 1.1986 | 0.0307 | 0.0666 | -1.1392 |
| Hid1 | 0.0022 | 0.0695 | -1.1501 | 0.0000 | 0.0003 | 1.2460 |
| Hltf | 0.0017 | 0.0641 | 1.3296 | 0.0110 | 0.0291 | -1.2412 |
| Hrh3 | 0.0002 | 0.0337 | -1.3263 | 0.0093 | 0.0252 | 1.1963 |
| Hrk | 0.0006 | 0.0442 | -1.2008 | 0.0000 | 0.0003 | 1.2665 |
| Hsd17b10 | 0.0009 | 0.0518 | -1.1970 | 0.0087 | 0.0240 | 1.1404 |
| Iars2 | 0.0049 | 0.0945 | 1.1757 | 0.0006 | 0.0027 | -1.2415 |
| Icam5 | 0.0008 | 0.0495 | -1.2012 | 0.0034 | 0.0110 | 1.1642 |
| Ift22 | 0.0008 | 0.0495 | -1.2215 | 0.0002 | 0.0013 | 1.2532 |
| Impad1 | 0.0004 | 0.0393 | 1.1762 | 0.0003 | 0.0016 | -1.1813 |
| Inafm1 | 0.0003 | 0.0353 | -1.3010 | 0.0047 | 0.0145 | 1.2085 |
| Insyn2a | 0.0008 | 0.0501 | 1.9775 | 0.0254 | 0.0571 | -1.5043 |
| Jpt1 | 0.0021 | 0.0685 | -1.2000 | 0.0003 | 0.0015 | 1.2603 |
| Jun | 0.0017 | 0.0644 | 1.2051 | 0.0002 | 0.0011 | -1.2706 |
| Kank1 | 0.0012 | 0.0554 | -1.3872 | 0.0084 | 0.0234 | 1.2802 |
| Kansl1 | 0.0014 | 0.0590 | 1.2247 | 0.0001 | 0.0008 | -1.3023 |
| Kcnip2 | 0.0018 | 0.0647 | -1.3174 | 0.0007 | 0.0030 | 1.3655 |
| Kcnt1 | 0.0005 | 0.0406 | -1.3363 | 0.0001 | 0.0006 | 1.4161 |
| Kctd17 | 0.0018 | 0.0656 | -1.1942 | 0.0003 | 0.0017 | 1.2400 |
| Kremen1 | 0.0011 | 0.0547 | 1.6762 | 0.0044 | 0.0138 | -1.5107 |
| Lemd2 | 0.0024 | 0.0711 | -1.2442 | 0.0043 | 0.0134 | 1.2263 |
| Lime1 | 0.0002 | 0.0337 | -1.5589 | 0.0000 | 0.0003 | 1.7043 |
| Limk2 | 0.0045 | 0.0919 | -1.1598 | 0.0008 | 0.0036 | 1.2025 |
| Lnpep | 0.0001 | 0.0260 | 1.3255 | 0.0000 | 0.0001 | -1.4380 |
| Lrp10 | 0.0017 | 0.0631 | -1.3068 | 0.0136 | 0.0347 | 1.2136 |
| Lrp6 | 0.0030 | 0.0778 | 1.2361 | 0.0004 | 0.0018 | -1.3130 |
| Lrrc73 | 0.0030 | 0.0785 | -1.1909 | 0.0006 | 0.0027 | 1.2350 |

| | | | | | | |
|---|---|---|---|---|---|---|
| Lsm11 | 0.0045 | 0.0915 | -1.2902 | 0.0079 | 0.0222 | 1.2557 |
| Lzts3 | 0.0042 | 0.0883 | 2.2356 | 0.0063 | 0.0184 | -2.1492 |
| Man1b1 | 0.0047 | 0.0931 | -1.1579 | 0.0004 | 0.0019 | 1.2218 |
| Map3k12 | 0.0023 | 0.0704 | 1.4260 | 0.0010 | 0.0042 | -1.4834 |
| Masp1 | 0.0006 | 0.0442 | 1.4138 | 0.0022 | 0.0079 | -1.3428 |
| Med16 | 0.0013 | 0.0582 | -1.2261 | 0.0005 | 0.0023 | 1.2603 |
| Mef2a | 0.0009 | 0.0522 | 1.2458 | 0.0072 | 0.0206 | -1.1819 |
| Mei1 | 0.0012 | 0.0563 | -1.5484 | 0.0033 | 0.0107 | 1.4645 |
| Mgat4b | 0.0003 | 0.0363 | -1.2269 | 0.0157 | 0.0387 | 1.1291 |
| Mterf3 | 0.0026 | 0.0734 | 1.6014 | 0.0107 | 0.0284 | -1.4539 |
| Mtss2 | 0.0029 | 0.0767 | -1.1654 | 0.0411 | 0.0842 | 1.1000 |
| Mx2 | 0.0001 | 0.0248 | -7.7500 | 0.0001 | 0.0007 | 8.2500 |
| Mxi1 | 0.0048 | 0.0933 | 1.1624 | 0.0040 | 0.0126 | -1.1668 |
| Myh14 | 0.0032 | 0.0795 | -1.2604 | 0.0011 | 0.0044 | 1.3047 |
| Myh7 | 0.0005 | 0.0421 | -1.6697 | 0.0203 | 0.0478 | 1.3844 |
| Mypop | 0.0006 | 0.0442 | -1.2132 | 0.0027 | 0.0092 | 1.1737 |
| Naga | 0.0023 | 0.0710 | 1.6240 | 0.0009 | 0.0038 | -1.7350 |
| Nap1l3 | 0.0050 | 0.0955 | 1.2268 | 0.0269 | 0.0599 | -1.1730 |
| Nap1l5 | 0.0002 | 0.0334 | 1.2828 | 0.0000 | 0.0003 | -1.3318 |
| Napepld | 0.0007 | 0.0468 | 1.3980 | 0.0140 | 0.0355 | -1.2394 |
| Nebl | 0.0002 | 0.0309 | 1.3023 | 0.0016 | 0.0060 | -1.2278 |
| Nectin2 | 0.0023 | 0.0704 | -1.4946 | 0.0007 | 0.0030 | 1.5761 |
| Nptxr | 0.0009 | 0.0505 | -1.2533 | 0.0001 | 0.0005 | 1.3388 |
| Nrep | 0.0024 | 0.0711 | -1.3848 | 0.0309 | 0.0669 | 1.2443 |
| Nthl1 | 0.0031 | 0.0794 | -2.8333 | 0.0059 | 0.0174 | 2.5833 |
| Olfm3 | 0.0012 | 0.0554 | 1.2244 | 0.0001 | 0.0004 | -1.3231 |
| Onecut2 | 0.0009 | 0.0517 | 1.4151 | 0.0225 | 0.0518 | -1.2256 |
| Otof | 0.0001 | 0.0260 | -1.7240 | 0.0002 | 0.0013 | 1.6617 |
| Pcdh7 | 0.0007 | 0.0468 | 1.1869 | 0.0057 | 0.0169 | -1.1404 |
| Pfdn4 | 0.0000 | 0.0165 | 1.3747 | 0.0000 | 0.0000 | -1.4777 |
| Polr2j | 0.0016 | 0.0629 | -1.2153 | 0.0030 | 0.0099 | 1.1958 |
| Polr2l | 0.0010 | 0.0525 | -1.1679 | 0.0466 | 0.0932 | 1.0860 |
| Pp2d1 | 0.0025 | 0.0720 | -1.8660 | 0.0397 | 0.0821 | 1.4845 |
| Ppargc1a | 0.0002 | 0.0333 | 1.2481 | 0.0045 | 0.0140 | -1.1659 |
| Ppfia4 | 0.0014 | 0.0593 | -1.2294 | 0.0001 | 0.0008 | 1.3084 |
| Ppm1k | 0.0020 | 0.0680 | 1.2203 | 0.0025 | 0.0087 | -1.2132 |
| Ppp1r1a | 0.0003 | 0.0353 | -1.2807 | 0.0183 | 0.0439 | 1.1555 |
| Ppp1r2-ps2 | 0.0046 | 0.0925 | 4.5000 | 0.0282 | 0.0622 | -3.2727 |
| Praf2 | 0.0024 | 0.0713 | -1.1472 | 0.0001 | 0.0007 | 1.2138 |
| Preb | 0.0033 | 0.0815 | -1.2141 | 0.0011 | 0.0046 | 1.2509 |

| | | | | | | |
|---|---|---|---|---|---|---|
| Prkcz | 0.0054 | 0.0986 | -1.1132 | 0.0002 | 0.0012 | 1.1720 |
| Prmt2 | 0.0000 | 0.0188 | -1.3195 | 0.0000 | 0.0002 | 1.3433 |
| Prmt8 | 0.0002 | 0.0297 | -1.2114 | 0.0000 | 0.0002 | 1.2638 |
| Prmt9 | 0.0049 | 0.0945 | -1.2855 | 0.0127 | 0.0328 | 1.2417 |
| Prps1 | 0.0051 | 0.0955 | -1.1626 | 0.0014 | 0.0055 | 1.1933 |
| Prrt1 | 0.0036 | 0.0832 | -1.2219 | 0.0236 | 0.0539 | 1.1531 |
| Ptk2b | 0.0001 | 0.0247 | -1.2792 | 0.0000 | 0.0002 | 1.3209 |
| Ptov1 | 0.0020 | 0.0684 | -1.1639 | 0.0032 | 0.0105 | 1.1545 |
| Ptprs | 0.0042 | 0.0883 | -1.1475 | 0.0001 | 0.0006 | 1.2420 |
| Ptrhd1 | 0.0027 | 0.0750 | -1.2611 | 0.0049 | 0.0151 | 1.2378 |
| Qars | 0.0054 | 0.0986 | -1.1733 | 0.0026 | 0.0089 | 1.1933 |
| Qdpr | 0.0000 | 0.0165 | -1.2097 | 0.0008 | 0.0034 | 1.1443 |
| Rab27b | 0.0007 | 0.0468 | 1.2381 | 0.0003 | 0.0016 | -1.2609 |
| Rab43 | 0.0024 | 0.0713 | -1.3169 | 0.0079 | 0.0221 | 1.2623 |
| Ramp3 | 0.0001 | 0.0246 | -1.6385 | 0.0003 | 0.0017 | 1.5509 |
| Rapgef3 | 0.0032 | 0.0795 | -1.2740 | 0.0076 | 0.0214 | 1.2306 |
| Rasal1 | 0.0014 | 0.0592 | -1.3461 | 0.0001 | 0.0007 | 1.4956 |
| Rasd1 | 0.0043 | 0.0903 | 2.5000 | 0.0010 | 0.0039 | -3.0769 |
| Retreg1 | 0.0039 | 0.0867 | 1.1998 | 0.0029 | 0.0099 | -1.2073 |
| Rgs17 | 0.0046 | 0.0926 | 1.1623 | 0.0223 | 0.0516 | -1.1201 |
| Rhpn1 | 0.0054 | 0.0986 | -1.5319 | 0.0305 | 0.0662 | 1.3617 |
| Rlbp1 | 0.0012 | 0.0548 | -1.2947 | 0.0053 | 0.0160 | 1.2316 |
| Rnd2 | 0.0001 | 0.0275 | -1.3227 | 0.0008 | 0.0033 | 1.2641 |
| Rnf26 | 0.0021 | 0.0685 | -1.2733 | 0.0010 | 0.0042 | 1.3004 |
| Rpf1 | 0.0043 | 0.0903 | 1.2932 | 0.0006 | 0.0026 | -1.4084 |
| Rpl36-ps12 | 0.0050 | 0.0954 | -1.2357 | 0.0199 | 0.0470 | 1.1771 |
| Rps15 | 0.0046 | 0.0925 | -1.1550 | 0.0022 | 0.0079 | 1.1734 |
| Rps29 | 0.0005 | 0.0438 | -1.1655 | 0.0001 | 0.0006 | 1.2055 |
| S100a1 | 0.0012 | 0.0563 | -1.2633 | 0.0000 | 0.0003 | 1.3997 |
| Sars2 | 0.0021 | 0.0685 | -1.4868 | 0.0011 | 0.0044 | 1.5238 |
| Sbsn | 0.0009 | 0.0505 | -1.3747 | 0.0002 | 0.0010 | 1.4629 |
| Scn3b | 0.0000 | 0.0165 | -1.3450 | 0.0005 | 0.0022 | 1.2482 |
| Sel1l | 0.0028 | 0.0757 | -1.1352 | 0.0008 | 0.0035 | 1.1583 |
| Selenoi | 0.0003 | 0.0337 | 1.7246 | 0.0001 | 0.0008 | -1.7796 |
| Senp3 | 0.0003 | 0.0353 | 2.2914 | 0.0000 | 0.0003 | -2.8954 |
| Sh2b2 | 0.0001 | 0.0260 | -1.4835 | 0.0000 | 0.0002 | 1.5861 |
| Sh3bgrl3 | 0.0006 | 0.0442 | -1.1840 | 0.0011 | 0.0045 | 1.1685 |
| Shisal1 | 0.0021 | 0.0685 | -1.2340 | 0.0005 | 0.0024 | 1.2866 |
| Slc15a4 | 0.0010 | 0.0527 | -1.2557 | 0.0008 | 0.0034 | 1.2658 |
| Slc25a1 | 0.0040 | 0.0871 | -1.2813 | 0.0027 | 0.0092 | 1.2888 |

| | | | | | | |
|---|---|---|---|---|---|---|
| Slc27a3 | 0.0035 | 0.0829 | -2.4444 | 0.0005 | 0.0025 | 3.0741 |
| Slc29a4 | 0.0006 | 0.0452 | -1.9024 | 0.0467 | 0.0934 | 1.4309 |
| Slc30a3 | 0.0028 | 0.0757 | -1.2628 | 0.0185 | 0.0444 | 1.1914 |
| Slc4a10 | 0.0006 | 0.0442 | 1.3181 | 0.0005 | 0.0022 | -1.3264 |
| Slc8a2 | 0.0001 | 0.0275 | -1.2284 | 0.0047 | 0.0145 | 1.1426 |
| Smarca5 | 0.0049 | 0.0941 | 1.1994 | 0.0011 | 0.0045 | -1.2550 |
| Smcr8 | 0.0016 | 0.0629 | 1.7530 | 0.0004 | 0.0020 | -1.9530 |
| Smim15 | 0.0015 | 0.0608 | 1.2805 | 0.0073 | 0.0207 | -1.2258 |
| Snhg14 | 0.0001 | 0.0247 | 1.3444 | 0.0009 | 0.0037 | -1.2551 |
| Snrpg | 0.0003 | 0.0353 | -1.2456 | 0.0001 | 0.0007 | 1.2797 |
| Sorcs1 | 0.0020 | 0.0684 | 1.3391 | 0.0003 | 0.0014 | -1.4471 |
| Sos1 | 0.0003 | 0.0337 | 1.2400 | 0.0021 | 0.0075 | -1.1842 |
| Sst | 0.0003 | 0.0353 | -1.2654 | 0.0015 | 0.0057 | 1.2139 |
| Stag2 | 0.0052 | 0.0963 | 1.1774 | 0.0071 | 0.0204 | -1.1705 |
| Steap2 | 0.0003 | 0.0352 | 1.5075 | 0.0184 | 0.0442 | -1.2657 |
| Stum | 0.0026 | 0.0728 | -1.1697 | 0.0191 | 0.0455 | 1.1216 |
| Stx5a | 0.0002 | 0.0309 | -1.2660 | 0.0000 | 0.0003 | 1.3128 |
| Synj1 | 0.0033 | 0.0810 | 1.1385 | 0.0142 | 0.0359 | -1.1085 |
| Tceal1 | 0.0001 | 0.0274 | 1.3438 | 0.0001 | 0.0008 | -1.3422 |
| Tead3 | 0.0023 | 0.0704 | 2.4444 | 0.0042 | 0.0134 | -2.2000 |
| Tenm2 | 0.0001 | 0.0224 | 1.2443 | 0.0019 | 0.0069 | -1.1619 |
| Terf2 | 0.0029 | 0.0767 | -1.1575 | 0.0008 | 0.0033 | 1.1870 |
| Thra | 0.0001 | 0.0275 | -1.2566 | 0.0001 | 0.0008 | 1.2621 |
| Tmem223 | 0.0040 | 0.0871 | -1.1492 | 0.0011 | 0.0046 | 1.1780 |
| Tpp2 | 0.0036 | 0.0836 | 1.2239 | 0.0001 | 0.0006 | -1.3719 |
| Trnp1 | 0.0023 | 0.0706 | -1.1534 | 0.0276 | 0.0611 | 1.0991 |
| Tspyl4 | 0.0000 | 0.0165 | 2.2985 | 0.0006 | 0.0027 | -1.8746 |
| Ttbk2 | 0.0048 | 0.0941 | 1.1807 | 0.0246 | 0.0557 | -1.1327 |
| Ttc39b | 0.0020 | 0.0684 | 1.2750 | 0.0008 | 0.0033 | -1.3053 |
| Ttll13 | 0.0046 | 0.0929 | -2.0909 | 0.0017 | 0.0065 | 2.3636 |
| Tysnd1 | 0.0022 | 0.0702 | -1.2546 | 0.0001 | 0.0009 | 1.3635 |
| Ubald1 | 0.0007 | 0.0472 | -1.1420 | 0.0014 | 0.0055 | 1.1295 |
| Usf3 | 0.0020 | 0.0684 | 1.3079 | 0.0220 | 0.0510 | -1.2118 |
| Usp19 | 0.0001 | 0.0278 | 1.4405 | 0.0000 | 0.0001 | -1.5833 |
| Vash2 | 0.0044 | 0.0908 | -1.7765 | 0.0252 | 0.0568 | 1.5176 |
| Wbp11 | 0.0014 | 0.0593 | 1.1837 | 0.0002 | 0.0011 | -1.2367 |
| Wdtc1 | 0.0035 | 0.0829 | -1.2351 | 0.0100 | 0.0268 | 1.1940 |
| Wnt10a | 0.0025 | 0.0719 | -1.2458 | 0.0006 | 0.0025 | 1.2988 |
| Wnt4 | 0.0049 | 0.0942 | -1.2486 | 0.0154 | 0.0383 | 1.2038 |
| Wscd1 | 0.0021 | 0.0690 | -1.1791 | 0.0019 | 0.0069 | 1.1818 |

| | | | | | | |
|---|---|---|---|---|---|---|
| Xiap | 0.0003 | 0.0344 | 1.2234 | 0.0000 | 0.0004 | -1.2713 |
| Xpo4 | 0.0001 | 0.0275 | 1.5451 | 0.0003 | 0.0017 | -1.4980 |
| Zbtb38 | 0.0022 | 0.0692 | 1.1480 | 0.0010 | 0.0041 | -1.1635 |
| Zdhhc17 | 0.0002 | 0.0309 | 1.3860 | 0.0006 | 0.0028 | -1.3369 |
| Zfp341 | 0.0052 | 0.0964 | -1.2649 | 0.0201 | 0.0474 | 1.2054 |
| Zfp780b | 0.0006 | 0.0442 | 1.3720 | 0.0002 | 0.0009 | -1.4380 |
| Zfp827 | 0.0034 | 0.0825 | 1.2463 | 0.0007 | 0.0030 | -1.3136 |
| Zfp933 | 0.0023 | 0.0706 | -1.2520 | 0.0277 | 0.0613 | 1.1599 |
| Zyx | 0.0001 | 0.0246 | -1.4269 | 0.0001 | 0.0006 | 1.4286 |

**Table 2.** GO cellular component analysis of the 283 DEGs normalized with vitamin D supplementation.

| Term Name | - log10 of adjusted p value | Intersections |
|---|---|---|
| Somatodendritic Compartment (GO:0036477) | 2.6657 | MX2, PRKCZ, LZTS3, GABRD, CDK5, RAPGEF3, CYP46A1, GNAS, ABITRAM, KREMEN1, GABRA4, CPNE5, SST, ARHGAP33, PTK2B, ARFGEF2, CHL1, BCAN, SLC4A10, COMT, GABBR1, PPARGC1A, GRM8, SLC8A2, CPNE6, ADD1, SOS1, CX3CR1, HRH3 |
| Synapse (GO:0045202) | 2.2634 | AP1S1, PTPRS, MX2, PRKCZ, CALY, LZTS3, GABRD, CDK5, RAPGEF3, CYP46A1, FKBP1A, PRRT1, DBI, GABRA4, CACNB4, ARHGAP33, ANK2, PTK2B, ARFGEF2, BCAN, SLC4A10, COMT, GABBR1, GRM8, SLC8A2, RAB27B, CAMKV, SLC30A3, ADD1, PPFIA4, SOS1, FCHO2, HRH3 |
| Dendritic Tree (GO:0097447) | 1.9579 | MX2, LZTS3, GABRD, CDK5, CYP46A1, GNAS, ABITRAM, GABRA4, ARHGAP33, PTK2B, ARFGEF2, CHL1, BCAN, SLC4A10, COMT, GABBR1, PPARGC1A, SLC8A2, CPNE6, ADD1, CX3CR1, HRH3 |
| Neuron Projection (GO:0043005) | 1.9482 | AP1S1, PTPRS, MX2, PRKCZ, CALY, LZTS3, GABRD, GUCY2E, CDK5, RAPGEF3, CYP46A1, GNAS, ABITRAM, FKBP1A, GABRA4, CPNE5, QDPR, ARHGAP33, MAP3K12, PTK2B, ARFGEF2, CHL1, BCAN, SLC4A10, MYH14, COMT, GABBR1, PPARGC1A, GRM8, SLC8A2, SLC30A3, CPNE6, ADD1, CX3CR1, HRH3 |
| Intracellular Anatomical Structure (GO:0005622) | 1.7435 | VASH2, NREP, AP1S1, PTPRS, TYSND1, ZBTB38, SENP3, SELENOI, MX2, CRIP2, SEL1L, PRKCZ, RPF1, CALY, PPM1K, HSD17B10, PPP1R1A, GM14399, CBS, PTOV1, HAUS2, GRK6, RETREG1, SH3BGRL3, GPBP1L1, THRA, ATN1, MEF2A, QARS, GUCY2E, SARS2, ZYX, KANK1, GPT, XPO4, RASAL1, ZFP827, TPP2, AQR, CAMK1D, CDK5, KANSL1, NTHL1, RLBP1, TTBK2, PFDN4, RAPGEF3, PRPS1, AZI2, PRMT9, NEBL, IMPAD1, CYP46A1, CPSF1, GNAS, SMARCA5, ABITRAM, SLC15A4, FKBP1A, JPT1, PRRT1, USP19, MTSS2, DBI, EGLN2, RPS15, CCDC116, WNT4, ABI3, G3BP2, STX5A, KCNIP2, SLC25A1, RPS29, LSM11, RAMP3, JUN, QDPR, FNDC5, GPSM1, SH2B2, STAG2, ARSA, CACNB4, GDF11, CDA, NAPEPLD, TERF2, CD34, ZDHHC17, CHID1, ARHGAP33, CLIC1, MAP3K12, LNPEP, WDTC1, |

| | | |
|---|---|---|
| | | CCDC3, DDX50, RNF26, MGAT4B, HLTF, ANK2, GRCC10, B3GALT2, RAB43, RHPN1, PTK2B, DCTPP1, MYPOP, KCTD17, ARFGEF2, GABPB2, ZFP341, MYH14, COMT, GABBR1, ANAPC13, TCEAL1, PPARGC1A, STEAP2, RND2, RASD1, IFT22, ZFP780B, HRK, SLC8A2, WBP11, RAB27B, LEMD2, SMCR8, ETS1, NAP1L5, CDK2AP2, DNLZ, MTERF3, SCN3B, APLN, SLC30A3, LIMK2, XIAP, GDAP1, PREB, DHRS1, CPNE6, CIC, MYH7, ANKRD13B, CCNB2, ADD1, TRNP1, IARS2, NAP1L3, GSTM2, COPS9, POLR2L, CHMP1B, ONECUT2, GCLC, SOS1, B4GALNT1, OLFM3, EIF2A, C2CD5, AGPAT4, FCHO2, MXI1, CBR1, MAN1B1, GSTP1, SNRPG, TSPYL4, NAGA, CX3CR1, CEBPG, GRINA, FAM20C, TMEM223, HID1, GLI3 |
| Dendrite (GO:0030425) | 1.5173 | MX2, LZTS3, GABRD, CDK5, CYP46A1, GNAS, ABITRAM, GABRA4, ARHGAP33, PTK2B, ARFGEF2, CHL1, BCAN, SLC4A10, COMT, GABBR1, PPARGC1A, SLC8A2, CPNE6, ADD1, HRH3 |
| Cell Projection (GO:0042995) | 1.5112 | AP1S1, PTPRS, MX2, PRKCZ, CALY, LZTS3, GABRD, GUCY2E, KANK1, CDK5, TTBK2, RAPGEF3, CYP46A1, GNAS, ABITRAM, FKBP1A, MTSS2, ABI3, GABRA4, CPNE5, QDPR, SH2B2, ARHGAP33, MAP3K12, PTK2B, ARFGEF2, CHL1, BCAN, SLC4A10, MYH14, COMT, GABBR1, PPARGC1A, RND2, IFT22, GRM8, SLC8A2, TTLL13, SLC30A3, CPNE6, ADD1, C2CD5, CBR1, CX3CR1, HRH3, GLI3 |
| Plasma Membrane Bounded Cell Projection (GO:0120025) | 1.3827 | AP1S1, PTPRS, MX2, PRKCZ, CALY, LZTS3, GABRD, GUCY2E, KANK1, CDK5, TTBK2, RAPGEF3, CYP46A1, GNAS, ABITRAM, FKBP1A, MTSS2, ABI3, GABRA4, CPNE5, QDPR, SH2B2, ARHGAP33, MAP3K12, PTK2B, ARFGEF2, CHL1, BCAN, SLC4A10, MYH14, COMT, GABBR1, PPARGC1A, IFT22, GRM8, SLC8A2, TTLL13, SLC30A3, CPNE6, ADD1, C2CD5, CBR1, CX3CR1, HRH3, GLI3 |

**Table 3.** DEGs in *Mecp2*+/- cortex, relative to *Mecp2*+/+, that are normalized with both vitamin D supplementation and deficiency.

| | ***Mecp2*+/- Suppl vs Ctrl** | | | ***Mecp2*+/- Defn vs Ctrl** | | |
|---|---|---|---|---|---|---|
| **Gene ID** | **P-value** | **FDR** | **Fold change** | **P-value** | **FDR** | **Fold change** |
| 4933432I09Rik | 0.0159 | 0.0391 | 2.2857 | 0.0085 | 0.0483 | 2.6429 |
| Abitram | 0.0006 | 0.0027 | 1.3570 | 0.0012 | 0.0135 | 1.3068 |
| Ank2 | 0.0004 | 0.0019 | -1.2556 | 0.0214 | 0.0884 | -1.1356 |
| Arsa | 0.0011 | 0.0043 | -1.5236 | 0.0185 | 0.0801 | -1.3233 |
| Atn1 | 0.0011 | 0.0043 | -2.3376 | 0.0054 | 0.0359 | -2.0805 |
| Bcl7a | 0.0077 | 0.0217 | -1.2732 | 0.0075 | 0.0447 | -1.2911 |
| Car11 | 0.0005 | 0.0023 | 1.2102 | 0.0001 | 0.0034 | 1.2134 |
| Cbs | 0.0000 | 0.0002 | 1.6022 | 0.0011 | 0.0130 | 1.3762 |
| Ccnb2 | 0.0003 | 0.0013 | 3.0000 | 0.0039 | 0.0293 | 2.3750 |
| Cda | 0.0015 | 0.0057 | 2.2041 | 0.0049 | 0.0338 | 1.7959 |
| Chpf2 | 0.0002 | 0.0013 | 1.2476 | 0.0071 | 0.0431 | 1.1507 |
| Cpne5 | 0.0004 | 0.0020 | 1.2484 | 0.0188 | 0.0810 | 1.1392 |
| Cul9 | 0.0000 | 0.0003 | 1.4929 | 0.0008 | 0.0105 | 1.3250 |
| Frmpd3 | 0.0024 | 0.0085 | -2.0719 | 0.0052 | 0.0349 | -1.8982 |
| Gdf11 | 0.0003 | 0.0016 | 1.3804 | 0.0012 | 0.0140 | 1.3043 |
| Gli3 | 0.0001 | 0.0007 | -1.8631 | 0.0010 | 0.0126 | -1.4985 |
| Gm10860 | 0.0118 | 0.0307 | 1.8235 | 0.0003 | 0.0064 | 2.0000 |
| Gm14399 | 0.0418 | 0.0854 | 1.6471 | 0.0136 | 0.0654 | 1.7647 |
| Gm16485 | 0.0036 | 0.0115 | -1.2697 | 0.0148 | 0.0692 | -1.1815 |
| Gm20712 | 0.0007 | 0.0029 | 1.3018 | 0.0233 | 0.0934 | 1.1674 |
| Gm50402 | 0.0055 | 0.0165 | 2.1250 | 0.0027 | 0.0230 | 2.1250 |
| Gpsm1 | 0.0015 | 0.0057 | 1.3916 | 0.0047 | 0.0330 | 1.3234 |
| Grk6 | 0.0001 | 0.0007 | 1.2679 | 0.0054 | 0.0358 | 1.1430 |
| H2-DMa | 0.0002 | 0.0012 | 1.5596 | 0.0009 | 0.0118 | 1.4015 |
| Jpt1 | 0.0003 | 0.0015 | 1.2603 | 0.0236 | 0.0940 | 1.1218 |
| Kank1 | 0.0084 | 0.0234 | 1.2802 | 0.0007 | 0.0095 | 1.3949 |
| Kcnt1 | 0.0001 | 0.0006 | 1.4161 | 0.0000 | 0.0014 | 1.4161 |
| Lemd2 | 0.0043 | 0.0134 | 1.2263 | 0.0118 | 0.0600 | 1.1573 |
| Lime1 | 0.0000 | 0.0003 | 1.7043 | 0.0000 | 0.0009 | 1.6441 |
| Lzts3 | 0.0063 | 0.0184 | -2.1492 | 0.0110 | 0.0572 | -2.0260 |
| Map3k12 | 0.0010 | 0.0042 | -1.4834 | 0.0057 | 0.0369 | -1.3263 |
| Mei1 | 0.0033 | 0.0107 | 1.4645 | 0.0101 | 0.0541 | 1.3935 |
| Mterf3 | 0.0107 | 0.0284 | -1.4539 | 0.0152 | 0.0704 | -1.4258 |
| Mypop | 0.0027 | 0.0092 | 1.1737 | 0.0014 | 0.0154 | 1.2087 |
| Olfm3 | 0.0001 | 0.0004 | -1.3231 | 0.0003 | 0.0059 | -1.2366 |

| | | | | | | |
|---|---|---|---|---|---|---|
| Otof | 0.0002 | 0.0013 | 1.6617 | 0.0011 | 0.0128 | 1.4896 |
| Pp2d1 | 0.0397 | 0.0821 | 1.4845 | 0.0003 | 0.0062 | 1.8763 |
| Rab43 | 0.0079 | 0.0221 | 1.2623 | 0.0049 | 0.0339 | 1.2623 |
| Rlbp1 | 0.0053 | 0.0160 | 1.2316 | 0.0152 | 0.0703 | 1.1965 |
| Rps29 | 0.0001 | 0.0006 | 1.2055 | 0.0039 | 0.0293 | 1.1245 |
| Sbsn | 0.0002 | 0.0010 | 1.4629 | 0.0002 | 0.0045 | 1.3988 |
| Selenoi | 0.0001 | 0.0008 | -1.7796 | 0.0082 | 0.0474 | -1.4263 |
| Senp3 | 0.0000 | 0.0003 | -2.8954 | 0.0037 | 0.0282 | -2.0353 |
| Sh2b2 | 0.0000 | 0.0002 | 1.5861 | 0.0002 | 0.0050 | 1.4872 |
| Shisal1 | 0.0005 | 0.0024 | 1.2866 | 0.0025 | 0.0220 | 1.1764 |
| Slc27a3 | 0.0005 | 0.0025 | 3.0741 | 0.0078 | 0.0458 | 2.2222 |
| Sorcs1 | 0.0003 | 0.0014 | -1.4471 | 0.0218 | 0.0894 | -1.1712 |
| Steap2 | 0.0184 | 0.0442 | -1.2657 | 0.0066 | 0.0411 | -1.3083 |
| Stx5a | 0.0000 | 0.0003 | 1.3128 | 0.0041 | 0.0300 | 1.1669 |
| Tspyl4 | 0.0006 | 0.0027 | -1.8746 | 0.0053 | 0.0353 | -1.7129 |
| Ttll13 | 0.0017 | 0.0065 | 2.3636 | 0.0093 | 0.0513 | 1.9545 |
| Usp19 | 0.0000 | 0.0001 | -1.5833 | 0.0003 | 0.0060 | -1.4335 |
| Wbp11 | 0.0002 | 0.0011 | -1.2367 | 0.0079 | 0.0462 | -1.1324 |
| Zyx | 0.0001 | 0.0006 | 1.4286 | 0.0011 | 0.0132 | 1.2581 |

**Table 4.** DEGs in *Mecp2*+/- cortex, relative to *Mecp2*+/+, that are normalized with vitamin D supplementation and even further dysregulated in the cortex of *Mecp2*+/- on deficient diet.

| | ***Ctrl Mecp2+/- vs Mecp2+/+*** | | | ***Mecp2*+/- Defn vs Ctrl** | | |
|---|---|---|---|---|---|---|
| **Gene ID** | **P-value** | **FDR** | **Fold change** | **P-value** | **FDR** | **Fold change** |
| Agpat4 | 0.0051 | 0.0845 | -1.1728 | 0.0121 | 0.0607 | -1.1466 |
| B4galnt1 | 0.0020 | 0.0575 | -1.2064 | 0.0000 | 0.0010 | -1.3686 |
| Cbr1 | 0.0032 | 0.0693 | -1.1298 | 0.0016 | 0.0164 | -1.1440 |
| Clptm1 | 0.0009 | 0.0405 | -1.1790 | 0.0054 | 0.0360 | -1.1364 |
| Cops9 | 0.0007 | 0.0362 | -1.1729 | 0.0023 | 0.0210 | -1.1468 |
| Cyp46a1 | 0.0001 | 0.0184 | -1.2368 | 0.0056 | 0.0369 | -1.1352 |
| Egln2 | 0.0016 | 0.0512 | -1.1486 | 0.0000 | 0.0002 | -1.3241 |
| Gabbr1 | 0.0047 | 0.0821 | -1.1223 | 0.0117 | 0.0597 | -1.1047 |
| Gclc | 0.0015 | 0.0490 | 1.2387 | 0.0003 | 0.0057 | 1.2983 |
| Gm10076 | 0.0012 | 0.0452 | -1.1690 | 0.0000 | 0.0016 | -1.2552 |
| Grina | 0.0009 | 0.0405 | -1.1810 | 0.0179 | 0.0788 | -1.1117 |
| H2aj | 0.0011 | 0.0438 | -1.2572 | 0.0013 | 0.0147 | -1.2562 |
| Hid1 | 0.0048 | 0.0827 | -1.1501 | 0.0207 | 0.0862 | -1.1130 |
| Hrh3 | 0.0001 | 0.0221 | -1.3263 | 0.0003 | 0.0061 | -1.2877 |
| Polr2l | 0.0011 | 0.0447 | -1.1679 | 0.0032 | 0.0256 | -1.1453 |
| Ptk2b | 0.0008 | 0.0390 | -1.2792 | 0.0154 | 0.0710 | -1.1721 |
| Rpl36-ps12 | 0.0039 | 0.0760 | -1.2357 | 0.0007 | 0.0099 | -1.2925 |
| Sh3bgrl3 | 0.0008 | 0.0390 | -1.1840 | 0.0000 | 0.0010 | -1.2900 |
| Slc30a3 | 0.0027 | 0.0635 | -1.2628 | 0.0018 | 0.0181 | -1.2938 |
| Slc8a2 | 0.0003 | 0.0277 | -1.2284 | 0.0003 | 0.0057 | -1.2320 |
| Snhg14 | 0.0000 | 0.0070 | 1.3444 | 0.0005 | 0.0077 | 1.2093 |
| Ubald1 | 0.0011 | 0.0442 | -1.1420 | 0.0005 | 0.0078 | -1.1572 |
| Wnt10a | 0.0031 | 0.0679 | -1.2458 | 0.0041 | 0.0301 | -1.2289 |
